# Characterizing habit learning in the human brain at the individual and group levels: a multi-modal MRI study

**DOI:** 10.1101/2022.12.12.520188

**Authors:** Rani Gera, Maya Bar Or, Ido Tavor, Dana Roll, Jeffrey Cockburn, Segev Barak, Elizabeth Tricomi, John P. O’Doherty, Tom Schonberg

## Abstract

The dual-process theory of action control postulates that there are two competitive and complementary mechanisms that control our behavior: a goal-directed system that executes deliberate actions, explicitly aimed toward a particular outcome, and a habitual system that autonomously execute well-learned actions, typically following an encounter with a previously associated cue. In line with dual-process theory, animal studies have provided convincing evidence for dissociable neural mechanisms, mainly manifested in cortico-striatal regions, involved in goal-directed and habitual action control. While substantial progress has been made in characterizing the neural mechanism underlying habit learning in animals, we still lack knowledge on how habits are formed and maintained in the human brain. Thus far only one study, conducted more than a decade ago by Tricomi et al. (2009), has been able to induce habitual behavior in humans via extensive training. This study also implicated the posterior putamen in the process, using functional magnetic resonance imaging (fMRI). However, recent attempts to replicate the behavioral results of this study were not successful. This leaves the research of human habits, and particularly the research of their formation through extensive repetition, as well as their neural basis, limited and far behind the animal research in the field. This motivated us to (1) attempt to replicate the behavioral and imaging main findings of Tricomi et al., (2) identify further functional and microstructural neural modifications associated with habit formation and manifestation, and (3) investigate the relationships between functional and structural plasticity and individual differences in habit expression. To this end, in this registered report we used Tricomi et al.’s free-operant task along with multi-modal MRI methods in a well-powered sample (n=123). In this task participants’ sensitivity to outcome devaluation (an index of goal-directed/habitual action control) is tested following either short or extensive training. In contrast to our hypothesis, we were not able to demonstrate habit formation as a function of training duration nor were we able to relate any functional or microstructural plasticity in the putamen with individual habit expression. We found that a pattern of increased activations in the left head of caudate that re-occurred across each day’s training is associated with goal directed behavior and that increased processing of devalued cues in low-level visual regions was indicative of goal-directed behavior (and vice versa). In a follow-up exploratory analysis comparing habitual and goal-directed subgroups within each experimental group, we found that elevated activations in frontoparietal regions during early stages of training, as well as increased reactivity towards still-valued cues in somatosensory and superior parietal regions were found in individuals that were more inclined to perform goal-directed behavior (compared with more habitual individuals). Taken together, regions commonly implicated in goal-directed behavior were most predictive of individual habit expression. Finally, we also found that differential patterns of training-related microstructural plasticity, as measured with diffusion MRI, in midbrain dopaminergic regions were associated with habit expression. This work provides new insights into the neural dynamics involved in individual habit formation/expression and encourages the development and testing of new, more sensitive, procedures for experimental habit induction in humans.

## Introduction

Action control, acquired through instrumental learning, is hypothesized to be determined by an interplay between two distinct systems^1,2^: one responsible for goal-directed behaviors and another that forms and executes habitual behaviors. Goal-directed behavior relies on a relatively careful consideration of the available information and is particularly dependent on the association between a response and its outcome (R-O)^2,3^. Therefore, it allows executing carefully planned actions and the flexibility to adjust them in order to maximize desired outcomes when circumstances change (e.g., ^4^). However, this behavior is cognitively taxing. In contrast, habitual behavior is considered to be automatic and is cognitively undemanding^2,5^. It relies on the association between a stimulus and a response (S-R) and thus relatively fixed and insensitive to changes in outcome value. Newly acquired instrumental actions are at first goal-directed and are sensitive to changes in outcome value. However, with extensive training, namely through behavioral repetition, a qualitative shift emerges as responding becomes autonomous and is automatically triggered by an associated stimulus, regardless of changes in outcome value or in R-O contingency^6,7^ (for review see^3^). Notably, the vast majority of the evidence supporting this dual-system account of action control and its dynamics has been accumulated from animal research^8^ (mostly rats), whereas it is yet to be well-established and characterized in humans.

Habitual action control is typically adaptive, liberating mental resources while automating usually beneficial actions. However, an imbalance with the goal-directed system in the “struggle” for action control and more specifically, overreliance on the habitual system, constitutes a key feature in several psychopathologies. Such malfunction has been implicated in addiction^9–13^, obsessive-compulsive disorder (OCD)^14–17^, schizophrenia^18^, autism^19^, social anxiety^19,20^, Tourette syndrome^21^ and obesity^22^. Thus, characterizing the interplay between these two systems and specifically the shift from goal-directed to habitual control, as well as understanding the underlying neural mechanisms, are of great importance for psychotherapeutical and clinical interventions.

A substantial landmark in the habit learning field has been the development of the reinforcer devaluation paradigm in rodents^2^. This paradigm successfully dissociates goal-directed from habitual behavior, based on whether a learned action had become insensitive to outcome devaluation (usually through food satiation or conditioned taste aversion). It became a pivotal measurement method of habits, yielding fruitful insights in animal research. A major contribution of this paradigm has been its capacity to demonstrate the transition from goal-directed behavior (sensitive to outcome devaluation) to habitual behavior (insensitive to outcome devaluation), as a result of extensive instrumental training^6^. Different human paradigms, aimed to utilize the devaluation sensitivity criterion, have been developed since (e.g. ^4,10,17,19,23–26^). The majority of these tasks aimed to distinguish S-R from R-O action control to point at inter-individual differences or pathologic group tendencies (e.g. ^10,17,19^), identify relevant modulators (e.g. ^24^), or discern their neural correlates (e.g. ^4,25,26^). However, apart from the one study conducted by Tricomi et al.^23^, habit learning, as acquired through behavioral repetition, has not been clearly demonstrated.

Neuroscientific research has identified distinct brain regions corresponding with the formation and execution of goal-directed behavior and habits. In rodents, the dorsomedial striatum (DMS), the prelimbic cortex (PL) and the nucleus accumbens are the main regions implicated in goal-directed control^27–31^. The execution of habitual responding is particularly dependent on the dorsolateral striatum (DLS)^32,33^, gradually gaining response-control across the course of learning^34^. In humans, homologous to the rodent DMS, the anterior caudate nucleus, and homologous to the PL, the ventromedial prefrontal cortex (vmPFC)^3^, were implicated in goal directed behavior^4,23,35–37^. In contrast, the research of the neural mechanisms underlying habitual control in humans is limited. There is evidence that the human putamen is homologous to the rodent DLS in habitual action control^23,36,37^. McNamee et al. implicated the posterior putamen in stimulus-triggered actions, suggesting it has a specific role in habit-associated S-R encoding^37^. De Wit et al. found that elevated gray matter density in the posterior putamen and white matter tract strength between this region and the premotor cortex are linked to individual tendency to habit-like performance^36^. However, thus far only Tricomi et al.^23^ implicated the posterior putamen in habit learning, using functional magnetic resonance imaging (fMRI).

Based on accumulated knowledge, demonstrating habit induction in healthy humans and as follows, characterizing its underlying neural mechanisms, is still a considerable challenge. Although it was extensively demonstrated in animals for decades, only the above-mentioned procedure by Tricomi et al.^23^ was able thus far to demonstrate the shift from goal-directed to habitual control through extensive training in humans and point at relevant neural mechanisms. However, these prominent findings have yet to be successfully replicated. Moreover, a recent study reported five failures in experimental habit induction^38^, two of which were attempts to replicate the behavioral findings of Tricomi et al.^23^. Thus, the reliability of the habit induction procedure utilized by Tricomi et al.^23^ is currently unclear. Furthermore, to date, there is no other paradigm that has been able to reliably induce habits through extensive training in humans. Consequently, the neural mechanisms underlying the formation (through behavioral repetition) and manifestation of habits have yet to be characterized.

The discrepancy in results across human research, the gap between animal and human literature on habits and the fact that habits are such a fundamental feature of human behavior, motivated us to establish the experimental induction of habit learning and characterize its underlying neural mechanisms in humans.

In the current work we aimed to replicate and expand the findings of Tricomi et al.^23^ with a well-powered sample using multi-modal MRI methods. We conducted a power analysis using the *fMRIPower* tool^39^ of the effect demonstrated by Tricomi et al.^23^ in the right putamen, as defined by the automated anatomical labeling (AAL) atlas^40^. This analysis yielded an n=61 for the extensive training group where the effect in the posterior putamen was observed. Therefore, we aimed to obtain data from 122 valid participants: 61 in each of two groups, differ in their training duration: short vs. extensive. Apart from the replication attempt of the Tricomi et al.^23^ findings, this well-powered sample also allowed us to test for further functional correlates with habit formation and manifestation (see Materials and methods). Additionally, we used diffusion tensor imaging (DTI) scans to probe micro-structural brain plasticity related to habit formation. We chose to use this method as in recent years it has been repeatedly shown that DTI indices can point at learning-induced neuroplasticity in gray matter regions^41–44^. Finally, we constructed a parametric index based on task performance, that measures the level of individual habit expression and correlated this measure with structural and functional measurements to identify neural determinants of habit formation. Our target sample size was four times larger than the one used by Tricomi et al.^23^ and thus had the potential to reliably identify relevant effects at both the individual and group levels that were not possible in the original study.

### Hypothesis

We hypothesized that extensive training will render responding habitual. The study conducted by Tricomi et al.^23^ was carried out in a considerably different environment than the unsuccessful replication attempts of its behavioral findings^38^. The original study was performed inside an MRI scanner, whereas the replication attempts were performed in common behavioral setting. Critically, such discrepancy with regard to the induction of habits, which heavily rely on the associations between responses and cues and contexts, may have led to different behavioral effects. Factors that may have potentially biased the behavioral effect as a function of the different environment include: (1) participants inside the MRI scanner may experience more stress which promotes habit formation^24^; (2) on the other hand, volunteers for MRI experiments may be self-selected to have low rates of stress and anxiety, allowing the manifestation of more goal-directed behavior following short training, thereby sharpening the differential effect of short and extensive training on action control; (3) The unusual and salient context of the MRI environment may impose S-R associations more robustly; (4) The loud noise in the MRI scanner may exploit some cognitive resources which may reduce the reliance on the goal-directed system (As evidenced by the effect of cognitive load on characteristics and strategies related to goal-directed action control^45,46^); (5) reduced arousal inside the MRI scanner may negatively affect goal directed action control. Taken together, we presumed that at least some of these factors enhance the formation and/or manifestation of habitual responding as a function of training duration. Thus, we expected to successfully replicate the behavioral findings of Tricomi et al.^23^.

Nonetheless, we considered the consequences of the possibility that the behavioral data would not support our hypothesis. This should not affect the individual level analyses we planned to perform; however, it is crucial for the interpretation of the group level analyses of the imaging data. Therefore, we stated that in case we would not observe the hypothesized behavioral effect, we will conduct an exploratory analysis, in which we will define two well-distinct subgroups for each experimental group. One subgroup will include participants who had expressed habitual responding and the other will include those who had not. The clustering will be based on a habit index calculated from the behavioral data (see in Individual differences in functional MRI for details on the generation of this index). Then, we will compare these subgroups within training conditions to identify functional and micro-structural differences.

We hypothesized that regions within the corticostriatal network will be implicated in habit formation and expression. For an elaborated depiction of the hypotheses and their corresponding confirmatory analyses of the neuroimaging data see Table 1. We expected that changes in activity and micro-structural plasticity in the putamen will be involved in habit learning while similar indices in the anterior caudate and vmPFC will be implicated in goal-directed action control. Nevertheless, the (goal-directed action control associated) R-O contingencies in our task are very easy to learn. Thus, it is highly likely that the anterior caudate and the vmPFC would not be employed differentially enough throughout the task to yield an effect in most of our planned analyses (see Table 1 and Data analysis). Therefore, the analysis of these regions is considered exploratory unless noted differently in Table 1.

**Table 1.** A mapping between hypotheses and confirmatory analyses of the neuroimaging data.

| # | Hypothesis | Confirmatory analysis / examined contrast | Neuroimaging modality | subtitle in the text |
| --- | --- | --- | --- | --- |
| **Within-group analysis (3-day group)** |  |  |  |  |
| 1 | The putamen increases its cue sensitivity after extensive training when habitual action control is expressed (replicating the finding of Tricomi et al.<sup>23</sup>). | [task onset – rest onset] of the last two vs. first two sessions of training: ROI analysis of the average signal in the putamen. | fMRI | Replicating the effect found by Tricomi et al.<sup>23</sup> in the posterior (right) putamen |
| 2 | The putamen increases its cue sensitivity after extensive training when habitual action control is expressed. | [task onset – rest onset] of the last two vs. first two sessions of training. | fMRI | Replicating the effect found by Tricomi et al.<sup>23</sup> in the posterior (right) putamen |
| 3 | The putamen gradually increases its cue sensitivity as training progresses. | Linear trend analysis on the contrast of [task onset – rest onset] by assigning increasing linear trend weights to the 12 training sessions according to their chronological order. | fMRI | Training duration induced-effects |
| 4 | The putamen increases its cue sensitivity after extensive training within each day. | [task onset – rest onset] of the last vs. first sessions of training, averaged across days. | fMRI | Within-day training duration induced-effects |
| 5 | The putamen gradually increases its cue-sensitivity as training progresses within each day. | Linear trend analysis on the contrast of [task onset – rest onset] by assigning increasing linear trend weights across the four within-day training sessions averaged across days. | fMRI | Within-day training duration induced-effects |
| **Between-group analysis** |  |  |  |  |
| 6 | Devaluation exerts larger changes in activations in the anterior caudate and vmPFC following short training compared to extensive training. | Compare groups (3-day group vs. 1-day group) for [post-devaluation difference – pre-devaluation difference] formed by first contrasting [valued snack onset – devalued snack onset] for pre-and post-devaluation separately. | fMRI | Effect of devaluation as a function of training duration |
| 7 | Extensive training induces micro-structural plasticity in the putamen compared to short training, expressed as a reduction in MD. | A voxel-wise mixed-design ANOVA with factors of time (before / after training) and group (1-day or 3-day) with participant as a random factor for extracted MD maps. | DTI | DTI analysis |
| **Individual differences analysis (Behavioral and Neuroimaging)** |  |  |  |  |
| 8 | The putamen activations, as hypothesized in points 2-5, and anterior caudate and vmPFC activations, as hypothesized in point 6, positively and negatively correlate with individual expression of habits, respectively. | Each hypothesis corresponding to points 2-6 was tested separately. For point 6 the analysis was done after collapsing the data across both groups. We used the individual maps generated in points 2-6 and correlate them with the individual behavioral habit index. | fMRI | Individual differences in functional MRI |
| 9 | Micro-structural changes in the putamen correlate with individual expression of habits, i.e., the larger the change (reduction in MD), the larger the habitual action control. | Correlation between the individual difference between extracted MD maps (before vs. after) and the individual behavioral habit index. We ran this analysis separately for each group. | DTI | Individual differences in micro-structural MRI |

In point 1 the ROI analysis is based on the automated anatomical labeling atlas, which was used for the power analysis of the effect demonstrated by Tricomi et al.<sup>23</sup> in the right putamen. For all other analyses we used small volume correction analysis using a mask based on the Harvard-Oxford atlas. All the within-group and between-group analyses require that the behavioral results demonstrate habitual responding as a function of training duration (when comparing sensitivity to outcome devaluation between groups) in order to relate their neuroimaging results to habits. Abbreviations: ROI, region of interest; vmPFC, ventromedial prefrontal cortex; fMRI, functional magnetic resonance imaging; DTI, diffusion tensor imaging; MD, mean diffusivity.

## Materials and methods

### Data Sharing

Registered report protocol preregistration is available at the Open Science Framework: https://osf.io/385dx. This protocol received in-principal acceptance on 21 June 2019. Behavioral data, analysis codes and task codes are available through the Github repository: https://github.com/ranigera/MultiModalMRI_Habits. The imaging data in Brain Imaging Data Structure (BIDS) format is available at OpenNeuro (https://openneuro.org/datasets/ds004299/versions/1.0.0) and unthresholded statistical maps are available at NeuroVault (https://neurovault.org/collections/13090).

### Participants

We aimed to collect a sample size of 122 valid participants, randomly assigned to two groups receiving short (1-day) or extensive (3-day) training, each with 61 participants. This number was based on a power analysis calculated using the fMRIPower tool^39^ of the effect demonstrated by Tricomi et al.^23^ in the right putamen (Figure 1), as defined by the automated anatomical labeling (AAL) atlas^40^, yielding n=61 for the extensive training group (on which the effect was originally demonstrated).

**Figure 1.**
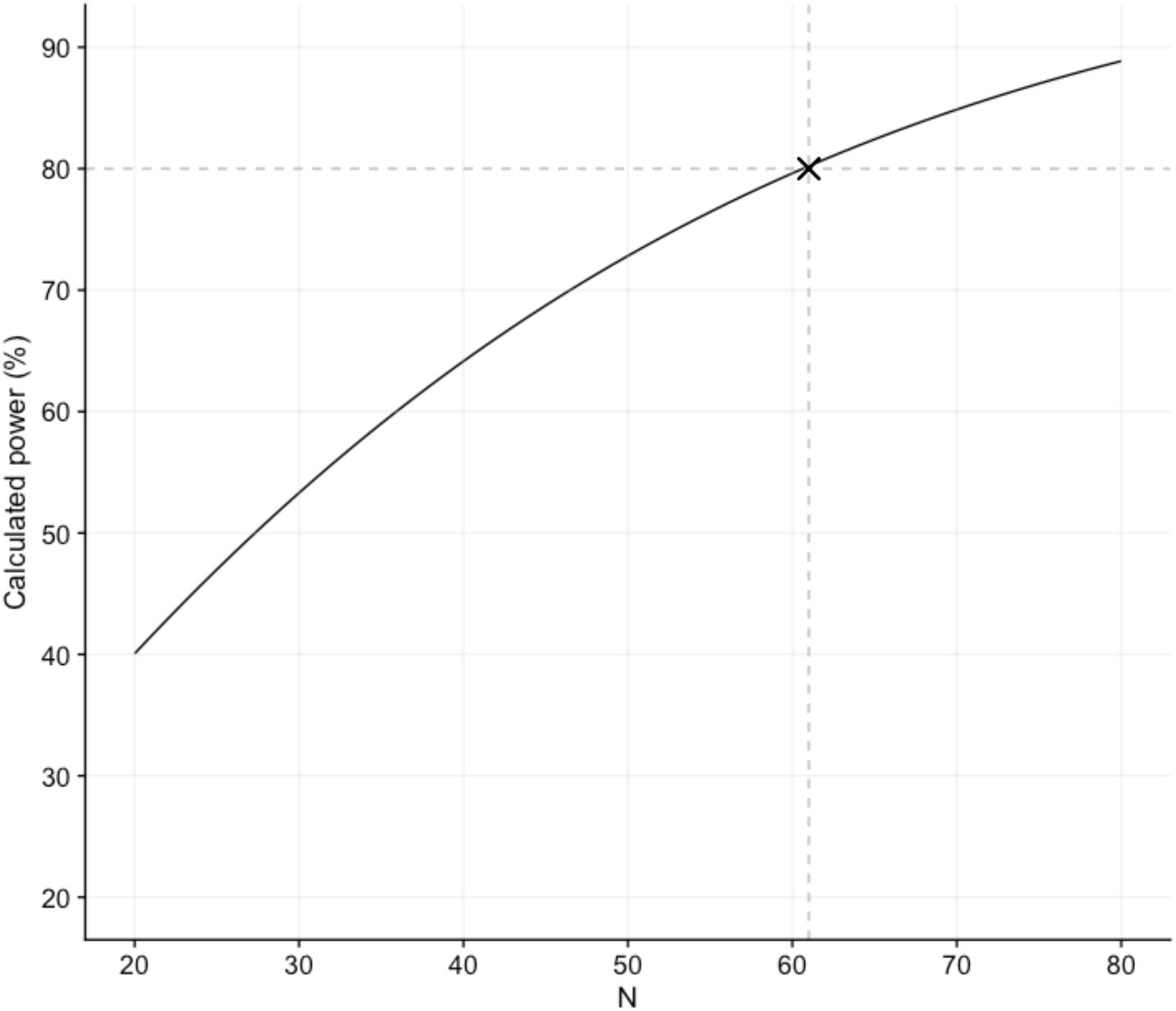
Results of the power analysis of the effect found by Tricomi et al.^23^ in the right putamen (as defined by the automated anatomical labeling atlas^40^), using the fMRIPower tool^39^. The horizontal dashed line represents estimated power of 80%. The vertical dashed line represents the number of participants required to cross the 80% power criterion (n=61). X represents where these lines meet.

To minimize a possible within-participant variability derived from diurnal variation, participants from the 3-day group were scheduled to participate at as similar time as possible at each day of the experiment. We confined participants to either morning or afternoon sessions. In addition, the experimenter collecting the data was the same person across all days of the experiment for each participant.

The study was approved by the institutional review board at the Sheba Tel Hashomer Medical Center and the ethics committee at Tel Aviv University. All procedures were performed in compliance with the relevant laws and institutional guidelines. We obtained informed consent from all participants prior to their participation in the experiment.

#### Recruitment

As food rewards were used in the experimental procedure (see Experimental procedure), participants were prescreened prior to their recruitment to ensure that they generally like eating snacks, do not restrict or limit their food consumption to avoid high calorie foods, are not vegan, do not suffer from food allergies in relation to snacks used in the experiment and are willing not to consume any food for 6 hours before arriving to each day of the experiment. Failing to comply with any of these criteria prevented participation in the experiment. They also rated their liking on a Likert pleasantness scale (ranging from -5, very unpleasant, to 5, very pleasant) toward three sweet and three savory snacks which were later used to choose from at the beginning of the experiment. To ensure participants’ desire to earn and eat snacks, participants who did not rate the highest sweet and highest savory snacks with at least +2 did not participate in the study. Finally, participants were asked to fill out the eating attitudes test (EAT-26^47^) to account for eating disorders. Participants with a score of 20 or above were excluded from participation in the study.

### Experimental procedure

Participants were scanned before, during and following a free-operant task (Figure 2), identical to the one used by Tricomi et al.^23^, aimed to render responding habitual as a function of extensive training. Upon their arrival and before scanning, participants were asked to taste three sweet and three savory snacks and choose their favorite one of each type. We used snacks comprised of small pieces: the sweet set included M&M, Skittles and a Maltesers-like Israeli snack of small chocolate balls; the savory set included potato chips, Doritos and cashews. The one sweet and one savory snacks chosen by each participant were then used throughout the entire experiment. Afterwards, participants entered the MRI scanner and first underwent DTI and resting-state scans. Then, they performed the free-operant task, consisting of three main stages: (1) training, (2) outcome devaluation and (3) extinction test. The training was comprised of 8-minute sessions. The amount of training sessions was varied between two experimental groups: either two sessions on a single day (1-day group) or 12 sessions, four on each of three consecutive days (3-day group). During all training sessions and the extinction test, participants were scanned using fMRI, whereas the devaluation procedure was conducted outside the scanner. Before starting and after completing the task phase on each day, participants’ DTI data was obtained. Anatomical scans were completed at the end of each day. An additional resting-state scan was performed after completing the task phase on the last day. Following the completion of the free-operant task and all scanning procedures, participants underwent a working memory capacity assessment and were asked to fill out a battery of questionnaires aimed at estimating the relationships between individual factors and tendencies to manifest habits as well as obtain self-report indices of habits. Participants also performed a variant of the two-step sequential decision-making task^48^, which has been shown to dissociate the use of model-free and model-based decision-making strategies. These strategies are hypothesized to reflect goal-directed and habitual action control. The experiment was programmed and run in Matlab (The MathWorks, Natick, Massachusetts, USA) using the Psychophysics toolbox^49^. For the working memory capacity assessment, we used a computerized task (see Working memory assessment) programmed and run using Inquisit Lab (Millisecond Software, Seattle, WA, USA).

**Figure 2.**
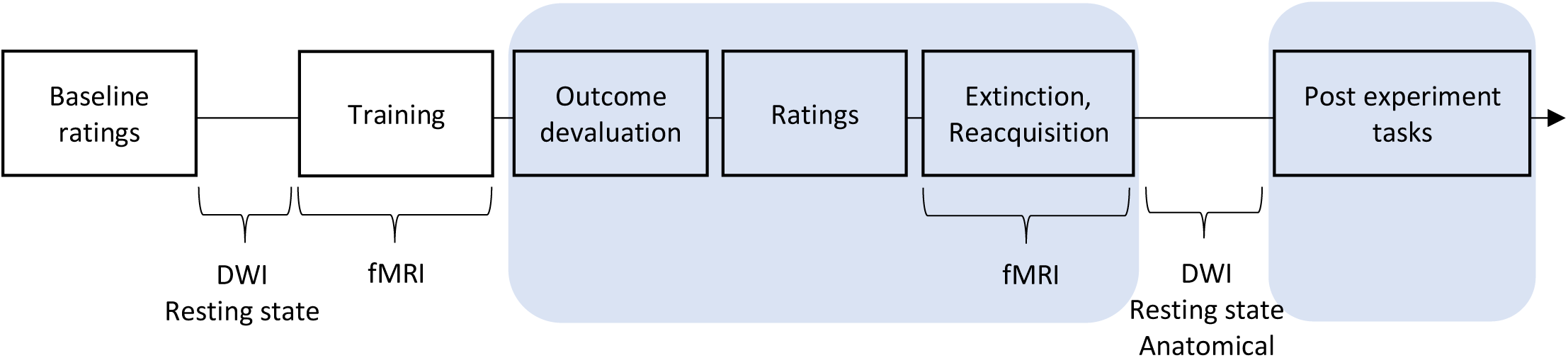
Procedure general outline. The behavioral tasks are presented along the arrow and below the line are the imaging scans. For the 3-day group the parts marked in blue were performed only on the third day. The other parts, including the magnetic resonance imaging (MRI) scans, were conducted on each of the three days, except for the resting-state scans, which were conducted only at the beginning of the first day and after completing the free-operant task on the last day. The 1-day group performed all stages on one day. Post experiment tasks included the administration of questionnaires, working memory assessment and a variant of the two-step sequential decision-making task^48^. Abbreviations: DWI, diffusion weighted imaging; fMRI, functional magnetic resonance imaging.

#### Free-operant task

##### Training

Each training session was comprised of 12 task blocks of 20 or 40 seconds and eight rest blocks of 20 seconds. During the task blocks, participants were trained on two Stimulus-Response-Outcome (S-R-O) associative structures to induce instrumental learning of two sets, each including a different discriminative stimulus. During each task block, participants were presented with a fractal (stimulus) along with an illustration of a corresponding button. Pressing this button (response) either yielded a snack reward (outcome), represented by its picture, to be consumed following the training, or a gray circle appearing briefly, indicating no reward. If any other button was pressed, the display did not change. Participants were instructed that they can press the button as often or as little as they like during the presentation of the fractal. We further instructed them that if they do not want any more of their favorite savory or sweet snack, they do not have to continue pressing and that otherwise they should try to earn as much reward as possible. Participants were also instructed to pay attention to the pairings of fractals and foods and that they will be tested on them later. Rewards were delivered on a 10-second variable interval (VI) schedule, implemented as 0.1 probability of reward becoming available every second. If a reward had become available, it was delivered on the next button press. Each fractal was associated with a particular button press and a particular snack consistently throughout the entire experiment for each participant, but counterbalanced across participants. A different fractal was used to indicate the rest blocks, where participants were asked not to respond. Each fractal appearance and disappearance marked the beginning and the end of a block, respectively. Block order was pseudo-randomized such that the same block type would not occur consecutively. At the beginning of the first day, before entering the scanner, participants were presented with the instructions for the training and underwent a short practice round. After completing each day’s training, participants received 0.5 snack pieces for their consumption for every snack reward won.

##### Outcome devaluation

Following completion of all training sessions, participants underwent a selective satiation^4,23,50,51^ outside the scanner, in order to devalue either the sweet or the savory snack (counterbalanced across participants). To accomplish that, participants received a large amount of one of the snacks, provided in a large bowl (∼1.4 liter), and were encouraged to consume it until it is no longer pleasant to them. Participants were further instructed that the size of the bowl is arbitrary and that they should ask for a refill if it has become empty and the snack is still pleasant to them. When deciding to stop eating, participants were asked if they are sure. The instructions accompanying this procedure were aimed to ensure that participants’ report of satiety is genuine and not due to the potentially intimidating laboratory setting.

##### Extinction

Subsequent to outcome devaluation, participants were placed back in the scanner, where they performed a three-minute extinction test. The extinction test was implemented in the same manner as the training phase and accordingly, participants were told that they will perform the same task as before. However, during this phase, responses were not rewarded. This phase consisted of nine 20-seconds blocks: three task blocks for each of the two associative structures and three rest blocks.

The goal of this part was to test whether responding has been rendered habitual, as measured by comparing the change in the response rate toward the valued outcome following devaluation with the change in response rate toward the devalued outcome following devaluation. A selective reduction in response rate toward the devalued outcome relative to the valued outcome serves as an evidence for goal-directed behavior, whereas the lack of such differentiation indicates insensitivity to outcome value and thus considered as an evidence for habitual control.

##### Reacquisition

We added a reacquisition test to exploratory evaluate the extent to which the selective satiation procedure is effective, and to examine how outcome presentation affects responding. Following extinction, participants performed another run comprised of nine 20-second blocks, three of each type, during which the outcomes were available again (as in the training stage). This phase was conducted inside the scanner to avoid context-related confounds.

##### Manipulation check

Participants were asked to rate on Likert scales their hunger (1, very full; 10, very hungry) and pleasantness towards each of the two snacks (-5, very unpleasant; 5, very pleasant) prior to each day’s training, following the devaluation procedure and at the end of the experiment upon completion of post-task scans. Regarding the pleasantness ratings, they were instructed not to rate the general pleasantness of each snack but rather to rate how pleasant a piece of each snack would have been pleasant to them at that moment. This allowed us to conduct a manipulation check for the outcome devaluation procedure, verifying it had indeed reduced the pleasantness for the devalued snack relative to the valued snack. Following devaluation, we used a similar pleasantness rating to obtain participants’ pleasantness toward the fractal images. Additionally, following the first day’s training, we assessed participants’ contingency awareness by presenting them each fractal and asking them to rate which of the snacks was more likely to be earned when pressing the corresponding button (-5, the specified sweet snack was more likely; 5, the specified savory snack was more likely) on a Likert-scale.

##### Working memory assessment

After completing the experiment, the participants were asked to perform a working memory capacity task. We used an automatic version of the Operation Span (OSPAN)^52^ procedure to assess working memory capacity for each participant. In this task, participants were asked to remember a series of letters while solving arithmetic problems. The score was the sum of items across all correctly recalled sets.

##### Two-step sequential decision-making task

Participants were asked to perform a variant of the two-step decision-making task^48^ designed to estimate the degree to which an individual uses a model-free or a model-based reinforcement learning strategy, while performing a multi-step learning and decision-making task. Participants were asked to choose between two actions, and according to their choice, they were stochastically transitioned into a visually identifiable outcome state where a reward (ranging from 0-100 points) is delivered following a key press. The task includes two key modifications to the original two-step task, each designed to improve task sensitivity to the action selection strategy used by individual participants. Firstly, participants were not asked to make a choice at the second level state; rather, they simply press a key to receive the reward associated with the outcome state. This design both simplifies the learning task from the perspective of the participant, and improves model identifiability by exacerbating the value discrepancy between first-level actions (see Kool et al.^53^ for a detailed discussion). Second, like the original two-step task, participants are asked to learn the probability of obtaining a reward at each outcome state; however, reward magnitudes will randomly vary from 1-100 point (0 being considered a loss). This modification exaggerates the learning signals associated with model-free (which learns according to observed reward magnitudes), and model-based (which learns about observed reward probability) controllers, further dissociating the behavioral signature associated with each strategy.

##### Questionnaires

After completing the experiment, participants were asked to fill out a battery of questionnaires aimed to explore potential moderators of behavioral and neural effects (or an interaction between them) observed in the experiment. The battery consisted of the following questionnaires: Obsessive Compulsive Inventory – Revised (OCI-R)^54^, Barratt Impulsivity Scale – (BIS-11)^55^, State-Trait Anxiety Inventory (STAI)^56^, Trier Inventory for Chronic Stress (TICS)^57^, A loss aversion questionnaire (using a modified version of a lottery choice task)^58,59^, Big Five Inventory (BFI)^60^ and Eysenck Personality Questionnaire – Revised (EPQ-R)^61^.

Additionally, we obtained individual reports of general tendency for habitual behavior using the Creature of Habits Scale (COHS)^62^ as well as individual sense of automaticity during the task using the Self-Report Behavioral Automaticity Index (SRBAI)^63^ with adjustments to fit our experiment (as done by Sanne de Wit et al.^38^). This allows us to explore the relationships between self-report indices of habits with both behavioral and neural indices.

Both questionnaires and working memory data were collected for exploratory analysis of the role of individual difference variables in habit formation and expression and to test the relationships between our task and self-reported habit indices. This analysis may serve to point at future study directions in the field and was analyzed as part of this work.

#### MRI data acquisition

We acquired imaging data using a 3T Siemens Prisma MRI scanner and a 64-channel head coil, at the Strauss imaging center located at Tel Aviv University. We acquired high-resolution T1-weighted structural images after each day’s task phase for anatomical localization using a magnetization prepared rapid gradient echo (MPRAGE) pulse sequence (Repetition time (TR) = 2.53 s, echo time (TE) = 2.99 ms, flip angle (FA) = 7°, field of view (FOV) = 224 × 224 × 176 mm, resolution = 1 × 1 × 1 mm). A DTI protocol included diffusion-weighted images acquired using spin-echo echo-planar-imaging pulse sequence (TR = 4 s; TE = 59.4 ms; Slice thickness=1.7 mm; image resolution 1.7 x 1.7) with a b value of 1000 s/mm2 in 64 nonorthogonal gradient directions. We also acquired five non-diffusion (b0) images and additional seven non-diffusion (b0) images with an opposite phase-encoding direction to account for susceptibility induced distortions. To accelerate acquisition, we used multiband acceleration factor (a simultaneous multi-slice method) of 2 and parallel imaging factor (iPAT) of 2.

For functional data we acquired T2*-weighted echo-planar images (TR = 1 s, TE = 30 ms, FA = 68°, FOV = 212 mm^2^, acquisition matrix of 106 × 106) with blood oxygen level-dependent (BOLD) contrast. Sixty-four interleaved axial slices, with their orientation tilted 30° from the anterior commissure-posterior commissure line to alleviate the frontal signal dropout^64^ were acquired. In-plane resolution was 2 × 2 mm, with a thickness of 2 mm and a gap of 0.4 mm to cover the entire brain. A multiband sequence^65^ with acceleration factor of 4 and iPAT of 2, in an interleaved fashion, was used. The same protocol with identical parameters was used to acquire six-hundred volumes of resting state data during a 10-minute scan, conducted prior to the first day’s beginning and following the last day’s completion of the free-operant task. During this scan participants were asked to rest and not to think about anything in particular while focusing on a fixation point. We used the eye-link 1000 plus eye-tracker to ensure their eyes are open.

### Data analysis

#### Exclusion criteria

Data from participants who did not complete all experimental parts, with the exceptions of the questionnaires, working memory assessment and two-step sequential decision-making tasks, was excluded. To guarantee that participants were indeed engaged in the task, i.e., wanted to earn the snacks and in a similar magnitude for both snacks, we averaged their pleasantness ratings that were obtained at the beginning of each day. An average of less than -1 or a difference of more than 3 points between the ratings of the sweet and savory snacks led to exclusion.

Additionally, to ensure similar amount of operant-training for both associative structures, data from participants with a difference of >2 SDs between the mean response rates toward the two snacks during the training procedure was excluded.

#### Behavioral analysis

Behavioral statistical analysis was carried out using R programming language (R Foundation for Statistical Computing, Vienna, Austria).

##### Manipulation checks

To verify that, as expected, the devaluation procedure selectively decreased pleasantness for the devalued snack we ran a 2 (phase: pre- or post-devaluation) x 2 (outcome: valued or devalued) repeated measures analysis of variance (ANOVA) on the pleasantness ratings. Additionally, we expected a reduction in hunger ratings following the devaluation procedure. To test this, we ran a paired-samples t-test to compare hunger ratings obtained pre- and post-devaluation.

##### Training duration induced-changes

We tested whether response rates were differentially affected by outcome devaluation and if the pattern is consistent with decreased sensitivity following extensive training. For each outcome type (valued and devalued) we calculated the difference between the average response rate during the extinction phase and the corresponding average response rate during the last training session. On this change measure we ran a mixed-model repeated measures ANOVA with a within-participant factor of Devaluation (valued or devalued outcome) and a between-participant factor of Group (1-day or 3-day) with participant as a random factor. We expected this analysis to yield an interaction effect, driven by a smaller reduction in the relative responding towards the devalued outcome in the 3-day group compared to the 1-day group, indicating the emergence of habitual responding. Such a pattern constitutes a replication of the behavioral results observed in Tricomi et al.^23^ (Figure 3A). Subsequently, conducted post hoc t-tests to determine whether there are significant differences between specific conditions. We verified that there are no significant differences in response rates between groups, or between the two outcomes during the last training session. For each group we compared changes in response rates for the valued and devalued outcomes following devaluation and further, compared response rates for the valued and devalued outcomes during extinction.

**Figure 3.**
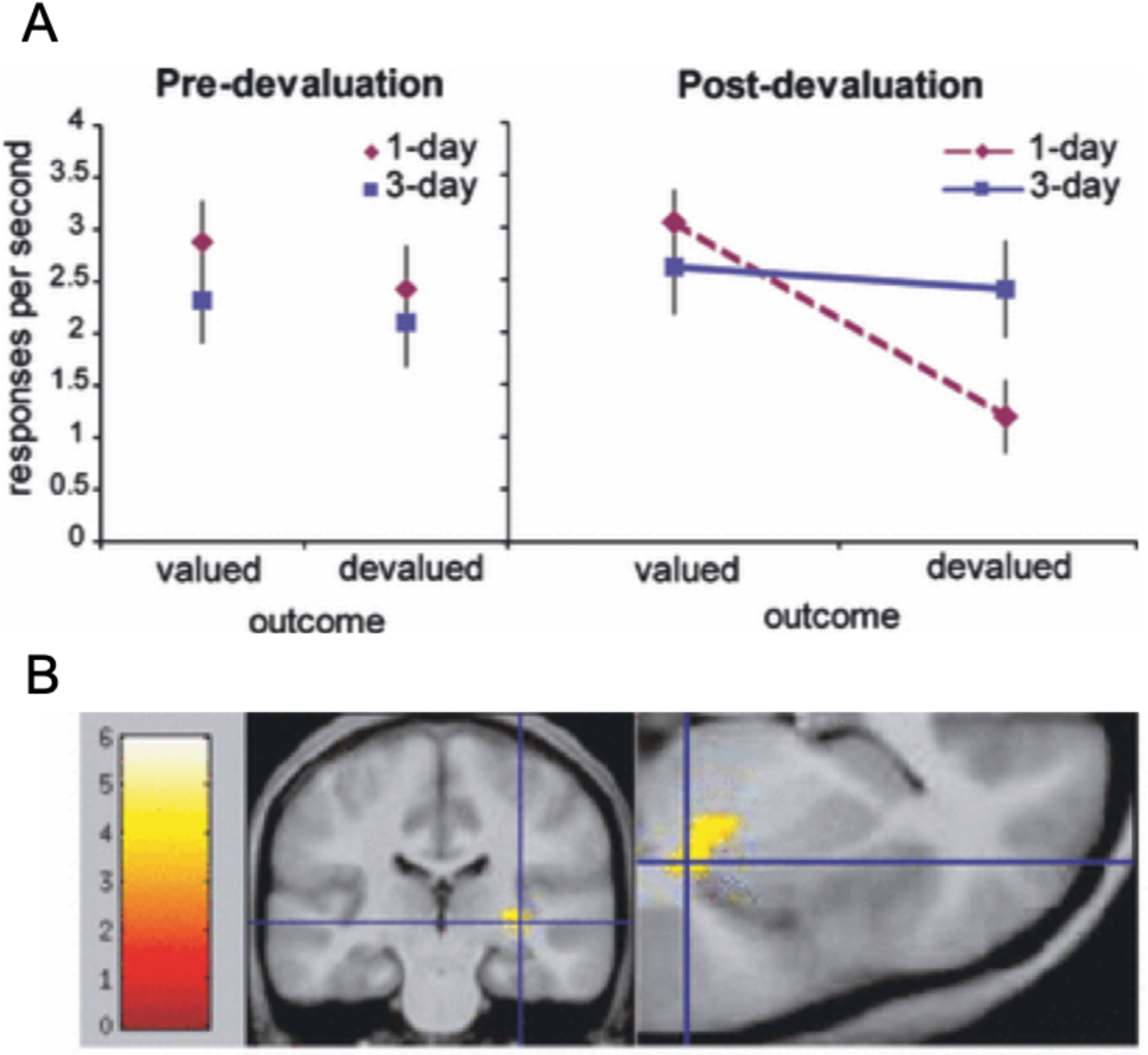
The main behavioral and neuroimaging results reported by Tricomi et al.^23^. **(A) Behavioral results.** During the last session of training, prior to the devaluation procedure (left), there were no significant differences in response rates between groups or when responding for the two food rewards (one which will be devalued through selective satiation and one which will not). During the extinction test following the devaluation procedure, response rates for the still-valued outcome remained high, as did response rates for the devalued outcome for the 3-day group. In contrast, response rates for the 1-day group for the devalued outcome were reduced, producing a significant training × devaluation interaction (*P* < 0.05). **(B) Neural correlates of habit learning**, as revealed by an increasing response with training to the onset of task blocks relative to the onset of rest blocks in the 3-day group. The right posterior putamen showed a significant increase in the [task onset – rest onset] contrast from the first two sessions to the final two sessions of training (*x* = 33, *y* = −24, *z* = 0; *P* < 0.001, *P*(cor) < 0.05). The blue crosshairs mark the voxels with the peak contrast value. * The figure and figure legend were adapted from Tricomi et al.^23^ with permission from the authors and the publisher (John Wiley and Sons).

#### Exploratory analysis of the two-step sequential decision-making task data

A computational model analysis of the two-step task data was planned to follow the mixture model approach outlined in Daw et al.^48^, in which model-free and model-based reinforcement learning agents are fit to the data according to a mixture parameter. Then, the extracted measure of individual degree of utilizing model-free/model-based strategies was planned to be used for exploratory analysis. We planned to examine its relationship with the individual habit index (as measured in the free-operant task) and with functional and microstructural indices we used for the individual differences analysis of the free-operant task data (see Neural correlates of individual tendency to form habits).

### MRI data analysis

#### fMRI data preprocessing

Raw imaging data in DICOM format was converted to NIfTI format and re-organized to fit to the Brain Imaging Data Structure (BIDS)^66^. We conducted preprocessing using fMRIprep^67–69^, which included: correction of each T1 weighted volume for bias field using N4BiasFieldCorrection^70^ and skull stripping using antsBrainExtraction (using OASIS template); Estimation of cortical surface using FreeSurfer^71^; Co-registration of the skullstripped T1w volume to skullstripped ICBM 152 Nonlinear Asymmetrical template^72^ using nonlinear transformation implemented in ANTs^73^; Functional data underwent motion correction using MCFLIRT^74^, then co-registration to its corresponding T1w volume using boundary-based registration with nine degrees of freedom – implemented in FreeSurfer^75^; Motion correcting transformations, T1 weighted transformation and Montreal Neurological Institute (MNI) template warp applied in a single step using antsApplyTransformations with Lanczos interpolation; Extraction of three tissue classes from T1w images using FSL FAST^76^; Voxels from cerebrospinal fluid and white matter were used to create a mask subsequently used to extract physiological noise regressors using CompCor^77^; Masks were eroded and limited to subcortical regions to limit overlap with gray matter and six principal components were estimated; Framewise displacement^78^ was calculated for each functional run using Nipype implementation.

We created confound files (tsv format) for each scan (each run of each task of each session of each participant), with the following columns: standard deviation of the root mean squared (RMS) intensity difference from one volume to the next (DVARS), six anatomical component based noise correction method (aCompCor)^77^, Frame-wise displacement^78^ (FD), and six motion parameters (translation and rotation each in 3 directions) as well as their squared and temporal derivatives (Friston 24-parameter model^79^). We added a single time point regressor (a single additional column) for each volume with FD value larger than 0.9, in order to model out volumes with extensive motion. Scans with more than 15% scrubbed volumes were excluded from analysis.

Following data processing by fMRIprep, for all other preprocessing procedures, we used FSL default settings, including prewhitening, high-pass filtering (using a 100-second cutoff) and spatial smoothing (using a 5 mm full-width-half-maximum Gaussian kernel).

#### fMRI analysis

We used FEAT (fMRI Expert Analysis Tool) of FSL^80^ for all fMRI analyses.

We conducted a general linear model (GLM) analysis, in which we planned to include for each participant the following regressors for each training session: ‘valued snack onset’, ‘devalued snack onset’ and ‘rest onset’ to model the onsets of the corresponding blocks, and ‘valued snack’, ‘devalued snack’ and ‘rest’ to model the entire corresponding blocks (see a slight necessary deviation from this plan in the fMRI main results). The regressors were convolved with a canonical double-gamma hemodynamic response function and their temporal derivatives were entered to the model. Additionally, the confounds produced by fMRIPrep described above were included in the model. This design matrix was regressed against each training and extinction session to generate parameter estimates for each participant for each session in the first-level analysis. For some of the analyses we first pooled together the BOLD data of ‘valued snack onset’ and ‘devalued snack onset’ to create a ‘task onset’ regressor (coinciding with Tricomi et al.^23^).

##### Replicating the effect found by Tricomi et al.^23^ in the posterior (right) putamen

To test for increased activation in the putamen as a function of training duration and corresponding to training duration-induced transition to habitual responding, we used the training BOLD data of the 3-day group. At the second-level (fixed effects), we averaged both task and rest onsets across the first two and last two sessions of training separately. Then, we tested a contrast of [task onset – rest onset] for the last two sessions of training compared with the first two. All second-level analyses across all participants of the 3-day group were used for a group analysis (mixed effects). The power analysis in fMRIPower is calculated to obtain an average effect across a specific region of interest (ROI) and in this case we thus extracted the average signal from the right putamen and tested for the replication of the effect from Tricomi et al.^23^ (Figure 3B).

In addition, we conducted a similar analysis, in which we performed small volume correction (SVC) analysis (see below for details) to test for the same contrast [task onset – rest onset] for the last two sessions of training compared with the first two in the (bilateral) putamen, anterior caudate and vmPFC.

##### Training duration induced-effects

We used the same first-level design matrix as described above to test for positive and negative correlations between the BOLD signal and training duration to identify changes in BOLD activity related to instrumental training as training proceeds. This was aimed to test shifts in balance of activity in specific regions of interest (putamen, anterior caudate and vmPFC). Similar to the analysis described above, we analyzed the BOLD training data for the training within the 3-day group. In the second-level analysis we used a linear trend analysis on the contrast of [task onset – rest onset] across the 12 training sessions, by assigning them linear weights [-13, -11, -7, -5, -3, -1, 1, 3, 5, 7, 11, 13] with respect to their chronological order. This second-level analyses were then used for a group analysis of the positive and negative correlations between our contrast of interest and training duration.

##### Within-day training duration induced-effects

Based on the within-day effect found by Tricomi et al.^23^ in the putamen and evidence for the role of context in the manifestation of habits^81^, we expected that habits will be re-formed on the slightly different contexts different days constitute, presumably slowly becoming more stable and less susceptible to changes in context. Thus, to test for within-day training effects, we first contrasted [task onset – rest onset] for each participant for each training session and then averaged each of the four sessions (separately) across the three days of training in the second-level (fixed effects) analysis. Then, we conducted the following analyses: (1) examined the contrast of [last (fourth) session – first session] in a group analysis; (2) constructed a linear trend analysis across the four training sessions by assigning them linear weights [-3, -1, 1, 3] with respect to their within-day chronological order during the second-level analysis. Then, we used a group analysis to test for positive and negative correlations between the BOLD signal and within-day training duration.

##### Effect of devaluation as a function of training duration

We tested for differential effects of training duration on relative changes in neural correlates toward valued and devalued outcomes following outcome devaluation. In a second-level analysis (fixed effects), we used a contrast of [valued snack onset – devalued snack onset] in the last training session (before the devaluation procedure) and the same contrast for the extinction test for each participant to produce estimates for pre- and post-devaluation difference, respectively. Then we used a contrast of [post-devaluation difference – pre-devaluation difference] to obtain a single map per participant representing the change in difference between the BOLD response to the valued snack block onsets vs. the devalued snack block onsets. All second-level analyses across all participants of both groups were used for a group analysis (mixed effects) where we compared extensive training (3-day group) with short training (1-day group).

We report group level whole-brain statistical maps thresholded at Z > 3.1 and cluster-based Gaussian Random Field corrected for multiple comparisons with a (corrected) cluster significance threshold of p= 0.05^82^.

#### Small volume correction (SVC) analysis

Based on prior research in animals and humans we hypothesized that goal-directed and habitual action control, as well as the shift between these two strategies, are substantially associated with the following brain regions: putamen, anterior caudate and vmPFC (see^3^ for review). Therefore, in addition to the whole-brain analyses, we conducted group level analyses for each of these regions, with a mask based on the Harvard-Oxford atlas, containing the voxels which are part of the region. Nevertheless, note that as pointed above, due to the nature of our task, we consider most of the anterior caudate and vmPFC SVC analyses exploratory (see Hypothesis and Table 1). The masks are shared with the data on OpenNeuro.

#### Secondary parallel fMRI analysis

In our planned fMRI data analysis presented above, we planned to focus on blocks’ onsets rather than the entire blocks. This was the case from several reasons: (1) comply with Tricomi et al. ^23^, (2) the entire task blocks include several components such as motor pressing which could hinder effects of interest, (3) fMRI may not be ideal in capturing such sustained effects and (4) habits are often conceptualized as an automated behavior initiated by the appearance of a cue, which fits well with the phasic event of cue presentation. However, behavior should be constantly under habitual control during the task blocks after a habit has been formed. Thus, as we have a relatively large sample that could be sensitive enough to identify relevant effects despite noisy conditions, we also ran the same analyses described above on the entire blocks’ data. To this end we used the same exact first-level GLM as above, however with the duration of the entire block and not only its onset. We used the ‘valued snack’, ‘devalued snack’ and ‘rest’ regressors instead of the ‘valued snack onset’, ‘devalued snack onset’ and ‘rest onset’, respectively. We consider this set of parallel group level analyses as secondary (exploratory) analyses and report only effects of interest that survived correction for multiple comparisons.

#### Exploratory resting-state fMRI analysis

We planned to use the acquired resting-state fMRI data for an exploratory analysis of the role of functional connectivity in habit formation and manifestation. We planned to generate seed to voxel connectivity maps based on our ROIs. Then, to conduct several analyses, including the following: comparing difference in changes in connectivity (before vs. after training) between groups, and examining separately in both groups how baseline connectivity predicts training outcome and how changes in functional connectivity (before vs. after training) correlate with training outcome. Special emphasis was planned to be placed on examining negative correlations between activity in the anterior caudate and the putamen and its relationship with behavioral manifestation of habits in both individual and group levels. We planned to report only effects of interest that survived correction for multiple comparisons.

#### DTI preprocessing

Pre-processing procedures of the diffusion images was conducted using FSL^80^. We used TOPUP^83^ to correct for susceptibility induced-distortions and EDDY^84^ to correct for motion and remove eddy currents artifacts. DTIFIT was used to fit a tensor model for each voxel and to extract mean diffusivity (MD) and fractional anisotropy (FA) maps. We planned to use FSL’s EPI_REG function for boundary-based registration^75^ of the diffusion maps (i.e., MD and FA) to structural data registration, followed by non-linear transformations to the MNI template using FNIRT^85^ (see a slight deviation from this plan in Notes on data collection and partial data exclusion).

#### DTI analysis

Our goal was to identify training duration-induced micro-structural plasticity, that as hypothesized, corresponds to a behavioral transition to habitual responding. All voxels of our ROIs (putamen, anterior caudate and vmPFC) were extracted according to the Harvard-Oxford atlas and a voxel-based mixed-design ANOVA with a within-participant factor of time (before or after training, namely, first and final scans) and a between-participant factor of group (1-day or 3-day) was planned to be performed with participant as a random factor. This analysis was planned to be carried out for both MD and FA measures. Yet, as evidence for MD task-induced changes is more robust, we consider the FA analysis as exploratory. We particularly expected significant micro-structural plasticity in the putamen as a function of training duration (manifested as an interaction between group and time). After performing this hypothesis-driven analysis, we also planned to conduct a whole-brain exploratory analysis. For all statistical tests of DTI data we used FSL’s Randomize tool^86^ to employ nonparametric permutation tests in which significance is assessed using cluster-based thresholding corrected for multiple comparisons based on cluster mass with a threshold of p<0.05.

#### Neural correlates of individual tendency to form habits

##### Individual differences in functional MRI

In addition to our group level analyses we aimed to identify the neural mechanisms underlying differences in individual tendencies to form and manifest habits. We constructed a parametric index, based on task performance, that measures the level of individual habit expression. For each participant we calculated the differences between the average response rate before devaluation (last training session) and after devaluation (during extinction) for the valued and the devalued outcomes. The difference between the two obtained measurements (valued change – devalued change) will be used as the habit index (with lower scores indicating more habitual behavior). We correlated this index with all individual functional data of the different indices described above, i.e., with the contrast of [task onset – rest onset] in the first two vs. the last two training sessions, training duration correlated activity, and a contrast of [post-devaluation difference – pre-devaluation difference]. The latter was analyzed after collapsing the data of both groups.

#### Individual differences in micro-structural MRI

To study the correlation of micro-structural changes with habit expression we performed a voxel-based correlation analysis between the individual habit index and the difference between the first and final scans (before and after training) of all voxels in our ROIs (putamen, anterior caudate and vmPFC). We analyzed both MD and FA measures (the analysis of the latter is considered exploratory) and used separate analysis for each group as they cannot be meaningfully compared.

## Main Results

In the sections below, we report all pre-registered analyses. We specifically note when presenting closely related exploratory analyses if integrated here (and not in the exploratory section) for the sake of consistency and fluency.

### Participants

In order to reach our target sample size of 122 valid participants (n=61 in each experimental group) we recruited 161 participants. Overall, 15 participants were excluded based on our pre-registered exclusion criteria: 13 participants were excluded due to a differential response rate (of >2 SD) towards the two snacks during the training phase, one participant was excluded due to low pleasantness rating for at least one of the snacks (average of less than -1) and another one for strongly preferring one of the snacks over the other (difference of >3), as measured before any snack consumption was made. Eleven participants were excluded because they did not complete the experiment, six due to technical reasons, four due to suspected brain abnormalities or medical findings and two were found out (post-recruitment) not be included within the scope of healthy participants. Our final sample included N=123 (61 participants in the short training group and 62 in the extensive training group), 61 females, aged 18 – 39 (mean = 24.7, SD = 3.81). Another participant’s fMRI data during the extinction test was not analyzed due to mistakenly acquiring it in 0° rather than 30° orientation from the anterior commissure - posterior commissure line, effectively omitting this participant’s only from the analyses of before vs. after devaluation.

### Notes on data collection and partial data exclusion

At around the middle of data collection we switched the specific type of Skittles we used (from the red to the green version) due to a shortage in stores. As we generalize across snacks and as participants chose their preferred sweet snack out of three options (at the beginning of the experiment) we assume this should not have affected our results.

The fMRI data from the second run in the first day of one participant was excluded due to excessive movement (more than 15% scrubbed volumes). We included this participant data in all analyses in which this run is first averaged with other runs by omitting it from in the averaging process, effectively excluding this participant only from the linear trend analysis across all days.

Fieldmap scans were not acquired for three participants in the extensive training group and thus we preprocessed their data without the fieldmap correction. Similarly, non-diffusion images of opposite phase-encoding direction were not acquired for five participants (four in the extensive training group and one in the short training group). Therefore, we preprocessed their DTI data without the susceptibility-induced distortion correction. In both cases, we tested whether the inclusion of these participants has a substantial influence on our findings by repeating all analyses without them. We found that removing these participants did not affect the pattern of any of our main fMRI and DTI results. Finally, two participants were omitted from the DTI analyses (both from the extensive training group) since some of their DTI data was not acquired.

Note that in order to achieve a satisfying registration for all diffusion data of all participants we had to slightly deviate from our initial preprocessing pipeline. Specifically, we ran FSL’s EPI_REG function on the first corrected B0 image rather than the MD and FA maps and then applied the native-to-MNI individual transformations formed by fmriprep.

#### Behavioral results

##### Training duration induced-changes

In contrast to our hypothesis we did not replicate the behavioral effects observed in Tricomi et al.^23^. Extensive training did not decrease sensitivity to outcome devaluation compared to short training as indicated by the lack of group x outcome type interaction effect on the change in response rate following outcome devaluation (F_1,121_ = 0.33, p = 0.567, ω^2^ = 0; Figure 5). This suggests that this procedure, at least with the currently used training parameters, may not be sensitive enough to either differentially induce or identify differences in habit formation as a function of training duration.

Although we did not find a group x outcome type interaction effect, we did find a main effect of outcome type (valued vs. devalued; F_1,121_ = 40.73, p < 0.001, ω^2^ = 0.07), driven by a smaller reduction in response rate for the still-valued outcome compared to the devalued outcome following outcome devaluation (t_60_ = 5.26 for the short training group, t_61_ = 3.88 for the extensive training group; p < 0.001). A comparison of the response-rate for the two outcomes strictly during the test phase (after devaluation) followed the same pattern (t_60_ = 5.13 for the short training group, t_61_ = 4.25 for the extensive training group; p < 0.001). We also verified that the difference between outcome types was not present prior to outcome devaluation (short training group: t_60_ = -0.87, p = 0.389; extensive training group: t_61_ = 0.72, p = 0.472). While these results are aligned with the interpretation that participants in both groups were goal-directed, this effect was mainly driven by a small subset of participants in each group, whereas the majority of participants (in both groups) responded habitually (See Exploratory subgroup analysis). Additionally, we found a main effect of group (F_1,121_ = 7.34, p = 0.008, ω^2^ = 0.05). This effect stems from a lower response rate in the short-training group compared to the extensive-training group prior to the outcome devaluation (t_120.3_ = -2.93, p = 0.004; see in Figure 5). This indicates a differential engagement pattern in the two training groups prior to the manipulation and could have potentially confounded the effects of training duration.

Exploratory: Following the work by Pool et al.^87^, who used the same procedure in behavior-only settings and found that the training duration effects on habit formation were moderated by the affective component of stress (composed of sub-factor such as anxiety, chronic worrying and social isolation), we conducted a similar analysis. A detailed description of the method and results of this analysis can be found in the Supplementary Material. Briefly, to adhere to the analysis used by Pool et al.^87^ we first conducted an exploratory factorial analysis on the relevant questionnaire data (STAI, TICS and BIS-11) to identify relevant factors and specifically to look for a factor that largely corresponds with stress affect (Supplementary Table 1). We then included the factor most corresponding with stress affect along with Devaluation (valued or devalued outcome), Group (1-day or 3-day) and all possible interactions as independent variables in a mixed-effects linear regression model where the dependent measure is the change in response rate following outcome devaluation. We used participant as a random factor. We found a trending interaction between the stress affect, devaluation and group (χ^2^_1_ = 3.55, p=0.059), indicating stress affect may modulate the interaction between devaluation and training length. We followed this analysis with a simple slope approach and tested the interaction between devaluation and group for participants low (-1 SD) and high (+1 SD) on the stress affect measure. We did not find an effect (though a weak trend) for participants with low levels of this measure (*β*= -0.38, 95%CI [-0.82, 0.06], *p*=0.097) and no significant effect for those with high levels (*β*= 0.24, 95%CI [-0.21, 0.68], *p*=0.304; Supplementary Figure 1). Thus, similar to Pool et al.^87^, but only descriptively, participants with low levels on the measure that corresponds with affective stress were trending towards habit formation only following extensive training whereas those with higher levels of this measure tended to respond habitually already after short training (Supplementary Figure 1).

##### Manipulation checks

The outcome devaluation procedure significantly reduced participants’ hunger levels (t_122_ = 14.16, p < 0.001; Figure 4) and differentially reduced participants’ pleasantness ratings toward the devalued snack vs. the still-valued snack as manifested in a significant phase x outcome interaction effect (F_1,122_ = 168.65, p < 0.001, ω^2^ = 0.25). This supports a successful induction of selective satiation by the outcome devaluation procedure.

**Figure 4.**
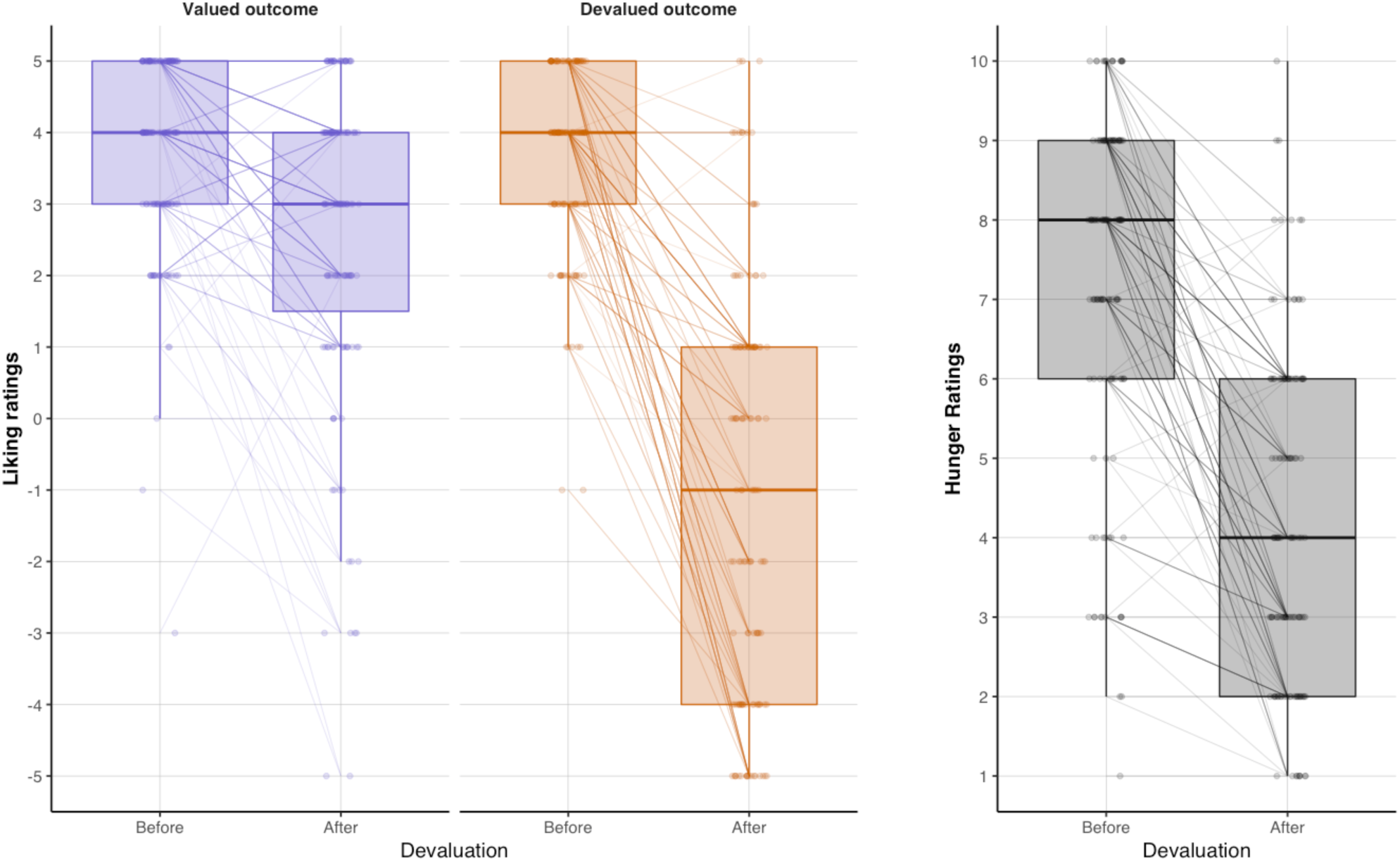
Manipulation checks of the outcome devaluation procedure. Liking ratings before and after outcome devaluation for the still-valued and devalued outcomes (left panel) and hunger ratings (right panel).

**Figure 5.**
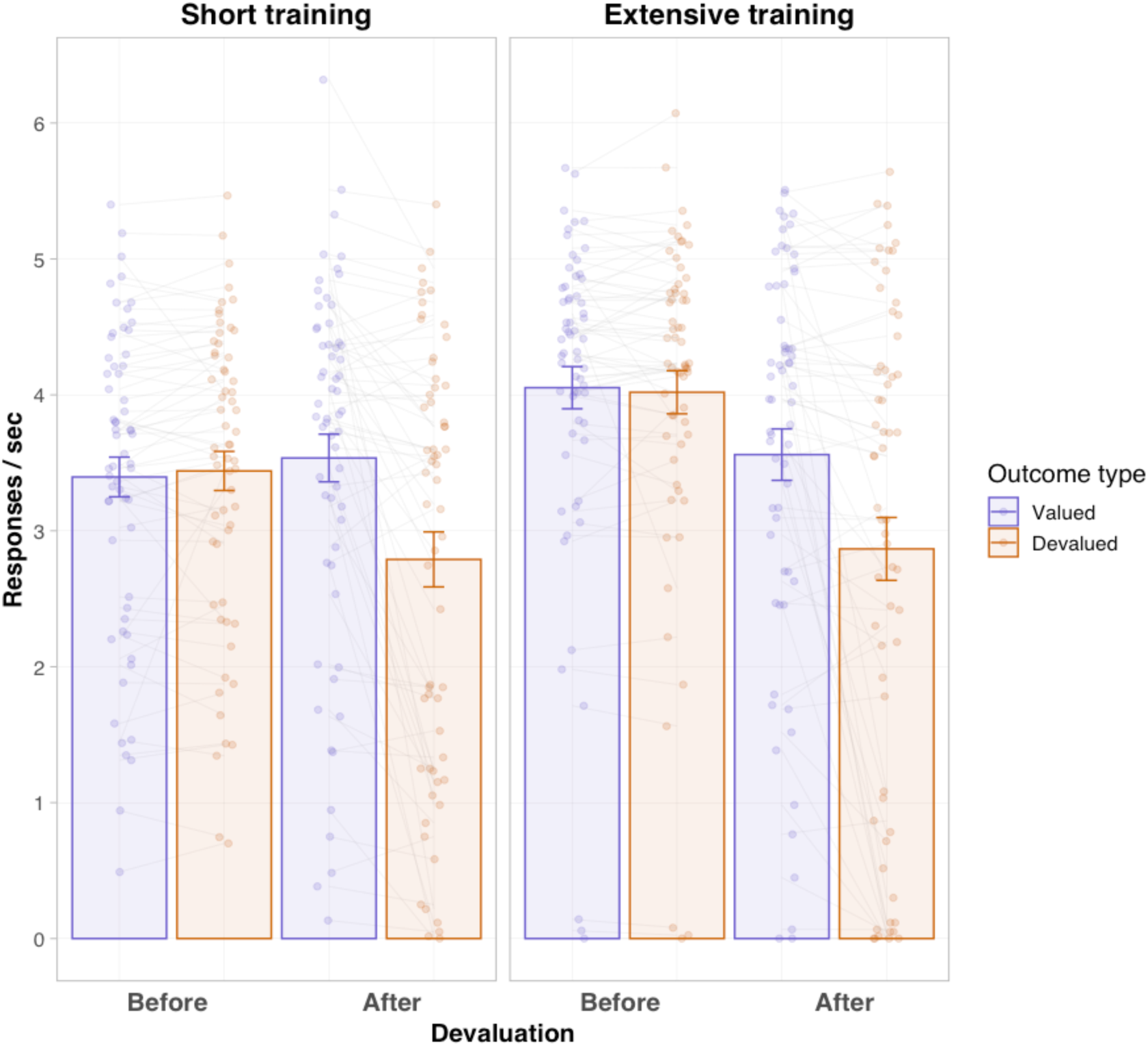
Response rates for the valued and devalued outcomes before and after devaluation in the short and extensive training groups.

Exploratory: We further tested whether these changes in liking and hunger were similar between groups and found a minor yet significant group x time interaction effect for the hunger ratings (F_1,121_=7.71, p < 0.006, ω^2^ = 0.02); see Supplementary Figure 2) but no difference for the liking ratings. The hunger effect was due to a slightly smaller decrease in hunger ratings in the short training group compared with the extensive training group, which may potentially indicate a slight difference in motivation between the two training groups.

### MRI results

As reported above, we did not replicate the differential effect of training duration on sensitivity to outcome devaluation as was found in Tricomi et al.^23^. Therefore, we only report the direct confirmatory pre-registered fMRI analyses in the putamen, corresponding to the main result of the original manuscript. We will present pre-registered individual-differences analyses that examined neural correlates of habit expression by correlating our neuroimaging data with the pre-registered individual habit index measure. In addition, we will report exploratory analysis testing relevant fMRI activation differences and differences in training-induced changes in MD and FA between goal-directed and habitual subgroups within each group, as identified by clustering the behavioral data within each training group (See Exploratory subgroup analysis below).

### fMRI main results

Below we report the fMRI results based on the block onset regressors. Originally, we pre-registered for these analyses a design matrix that also includes regressors for the entire blocks. However, this yielded high collinearity (due to the temporal derivative regressors) and therefore we separated them into two GLMs. The first analysis of the block onsets is reported in the pre-registered section and the additional analysis of the entire blocks was (as planned) added as an exploratory analysis and is reported below in the exploratory analyses section.

#### Direct replication attempt of the main fMRI results found in the putamen by Tricomi et al.^23^

##### ROI and SVC analyses

We conducted the ROI and SVC analyses to test changes in putamen sensitivity to task vs. rest cues following extensive training (last two vs. first two runs). This was done to provide a comprehensive report of our replication attempt of the results found by Tricomi et al.^23^ and to examine whether training duration leads to increased fMRI putamen reactivity to cues associated with action and rewards. Comparing the average change in activity in the putamen following extensive training ([task onset - rest onset] in the last two vs. first two runs) did not yield the hypothesized effect. This is not surprising as we did not obtain the behavioral results that would have allowed us to derive conclusions with respect to habit formation as a function of training duration. Surprisingly, the results were in the opposite direction, indicating a general decrease in activation (right putamen, t_61_ = -2.36, p=0.022; or when (exploratorily) testing the bilateral putamen, t_61_ = -2.60, p=0.012). Correspondingly, the SVC analysis in the putamen for the same contrast revealed a reduction in activation in the bilateral putamen (right putamen: cluster size=146, max Z-value=4.83, cluster-corrected p<0.001; left putamen: cluster size=94, max Z-value=4.16, cluster-corrected p<0.001; Figure 6). Nevertheless, see *Exploratory SVC analysis around the Tricomi et al. peak activation and visual inspection of raw activations across the putamen* (and Supplementary Figure 3) for a potential explanation of this surprising opposite than expected effect.

**Figure 6.**
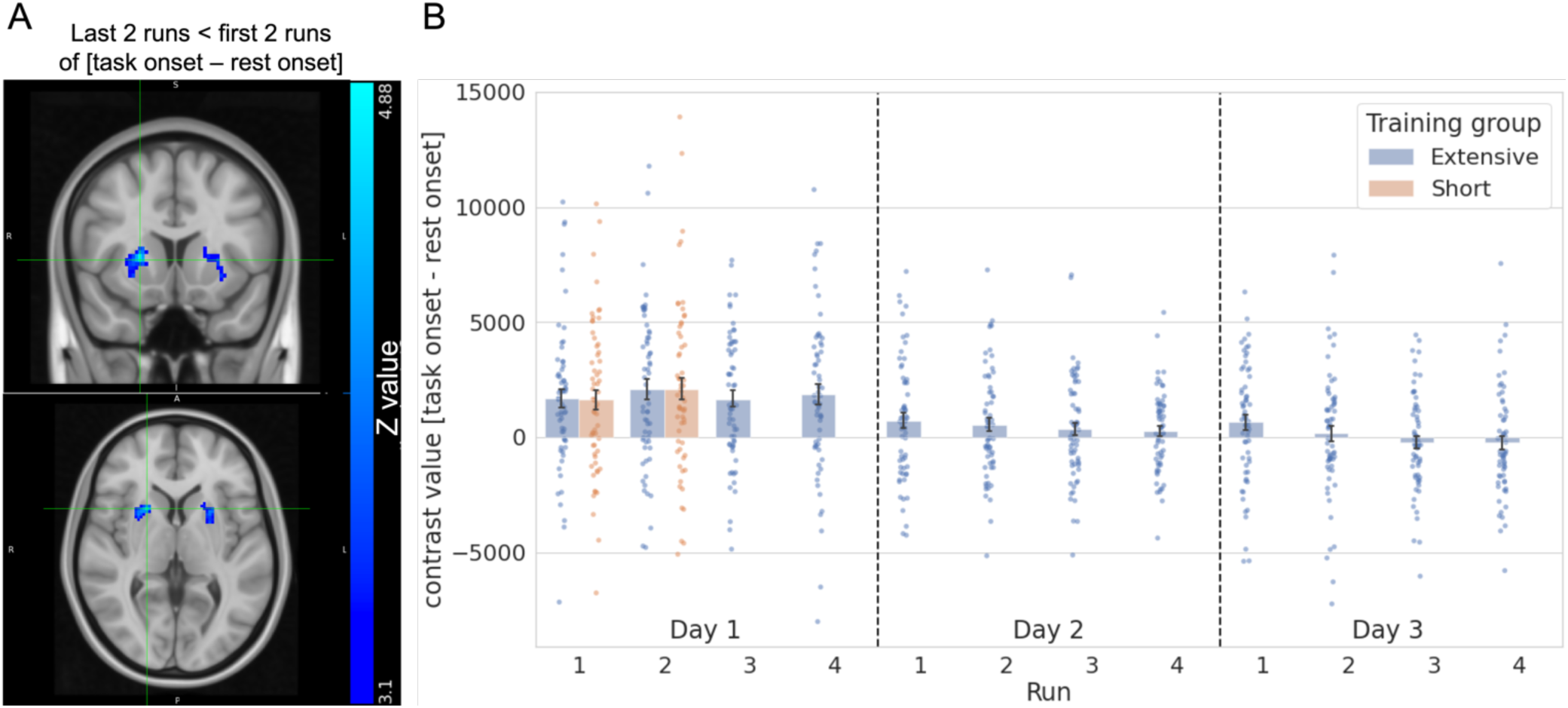
Replication attempt of the main fMRI results found in the putamen by Tricomi et al.^23^. **(A)** Results of small volume correction analysis in the bilateral putamen. The bilateral putamen reduced their sensitivity for task cues vs. rest cue following extensive training (last 2 runs vs. first 2 runs). Results were cluster corrected (p<0.05). **(B)** Plots are presented for illustrative purposes of the effects in panel A: averaged contrast values of task onset vs. rest onset within the identified significant clusters in the bilateral putamen (shown in panel A) in both groups. Error bars indicate 68% confidence intervals (equivalent to ±1 std error).

#### Individual differences in habit expression and related functional plasticity

##### Training duration-induced functional plasticity and habit expression

We did not find significant correlations in any of the analyses between the behavioral habit index and training duration-related changes in neural activity, neither when contrasting late vs. early stages of training nor when testing the presence of a linear trend throughout the entire training process in the extensive training group.

##### Within-day effects

Examining within-day effects, i.e., changes in cue sensitivity [task onset vs. rest onset] throughout daily training, did not reveal the pre-registered effect in the putamen. However, we did reveal in our exploratory yet pre-registered SVC analysis in the head of caudate a significant cluster in the left head of caudate both when contrasting the fourth vs. first (averaged) daily runs (cluster size=13, max Z-value=3.96, cluster-corrected p=0.028; Figure 7A) and when modelling a linear trend pattern across all (averaged) daily runs (cluster size=10, max Z-value=3.84, cluster-corrected p=0.048). The pattern of these results indicates that participants whose caudate maintained or reduced its reactivity to task cues tended to respond habitually (as revealed later following outcome devaluation), whereas participants tended to remain goal-directed when activations in the head of caudate became more sensitive to task cues as the daily training progressed (Figure 7B).

**Figure 7.**
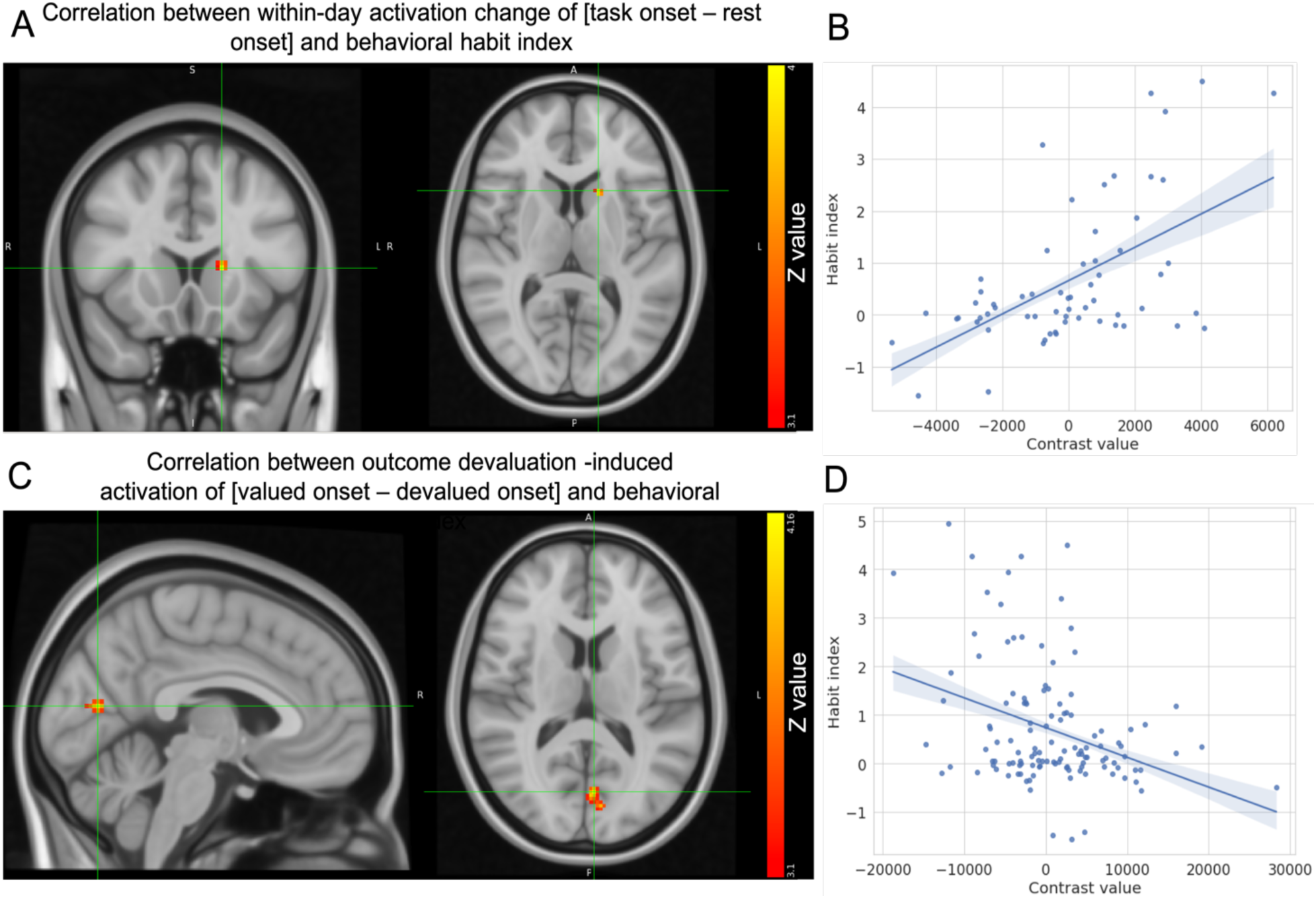
Habit expression related functional plasticity. (A) An SVC analysis in the head of caudate in the extensive training group of the correlation between within-day training-induced changes (averaged run 4 vs. averaged run 1 of [task blocks onset vs. rest block onsets]) and individual habit index revealed a cluster with a positive correlation in the head of the left caudate. An increase in activation in this region was associated with increased goal-directed behavior and vice versa. **(B)** Plots are presented for illustrative purposes of the effects in panel A: averaged contrast values within the significant cluster in the left head of caudate (shown in panel A). **(C)** Whole-brain analysis of the correlation between changes in BOLD response toward the valued vs. devalued cues following devaluation (after vs. before devaluation of [valued block onsets vs. devalued block onsets] and individual habit index revealed a negative correlation in a the left intracalcarine sulcus. Higher activation toward the devalued cue relative to the valued one were associated with goal directed behavior and vice versa. **(D)** Plots are presented for illustrative purposes of the effects in panel C: averaged contrast values within the significant cluster in the left intracalcarine sulcus (shown in panel C). Notes: (1) fMRI results were cluster-corrected (p<0.05). (2) The habit index is structured such that higher values indicate goal-directed behavior and values around 0 indicate habitual responding. (3) Shaded area around the fitted lines in the scatter plot indicates 68% confidence interval (equivalent to ±1 std error).

##### Neural corelates of differential value-based cue reactivity and habit expression

We tested whether changes in neural reactivity toward valued vs. devalued block onsets following outcome devaluation were associated with habit expression. We did not find significant correlations in our pre-registered ROIs. However, a whole-brain analysis across both groups revealed a significant negative correlation between goal-directedness and activity toward valued vs. devalued cues in the left intracalcarine sulcus (cluster size=43, max Z-value=4.12, cluster-corrected p=0.037; Figure 7C). As can be seen in the illustrative panel (Figure 7D) this effect in this low-level visual region, stems mainly from greater activation at the onset of the devalued cue relative to the still-valued cue in participants who exhibit goal-directed behavior.

We followed up this pre-registered analysis with an exploratory analysis to examine whether this effect was linked to the experimental group. To do so, we extracted the average contrast estimate (of [difference between valued and devalued after devaluation - difference between valued and devalued before devaluation]) for each participant in the identified cluster. We then ran a multiple linear regression with the habit index as the dependent variable and contrast value, group and their interaction as independent variables. We found no interaction effect (F_1,119_=1.54, *β*=0.21, 95%CI [- 0.13,0.55], p=0.217), implying the observed effect in the left intracalcarine sulcus is associated with habit expression regardless of training duration.

### DTI main results

The following details regarding the DTI analysis did not appear in the original pre-registration so are provided here for completeness. First, prior to running statistical analyses we spatially smoothed the MD and FA maps, with a 6 mm full-width-half-maximum Gaussian kernel. For the statistical analyses we set a voxel-based threshold of T > 3.1 for the MD and FA maps on which we employed the cluster mass threshold (of p<0.05).

#### Individual differences in habit expression and related micro-structural plasticity

We did not find any significant correlations between the behavioral habit index and training-induced changes in MD or FA in any of our ROIs and whole-brain analyses following either a short training or extensive training.

### Exploratory fMRI analyses

#### Exploratory SVC analysis around the Tricomi et al. peak activation and visual inspection of raw activations across the putamen

Intrigued by the discrepancy between the results obtained in our ROI analysis of the putamen and those obtained by Tricomi et al ^23^ we explored the possibility that our results were driven by activation changes in different sub-regions in the putamen than the sub-region identified in Tricomi et al.^23^. To this end we created a 5mm sphere mask centered around the peak MNI fMRI activation reported in Tricomi et al.^23^ and repeated the same procedure in an exploratory analysis. The results of this analysis descriptively indicate a trend in increase in bilateral putamen reactivity to learned cues following extensive training (t_61_ = 1.923, p=0.059). Moreover, a visual inspection of the unthresholded z-score maps of the same contrast reveal a clear differential response profile of the anterior and posterior subregions of the bilateral putamen, such that activation (to learned cues) decreases following extensive training in the anterior parts and appears to increase (though to a smaller extent) in the posterior parts (see Supplementary Figure 3). We consider these findings in the discussion section in the context of a proposal that distinct sub-regions in the putamen are responsible for these different increasing and decreasing effects as a function of training.

In addition, we repeated all of the planned individual differences and exploratory subgroup fMRI analyses (the latter are detailed below) using this focused sphere mask instead of the whole putamen mask to test whether our hypothesized effects would be revealed and captured when focusing on this specific region. We did not find any significant effect in any of these analyses thus preventing us from relating any activation patterns in this specific posterior putamen region (focused around the peak activation found by Tricomi et al ^23^) with sensitivity to outcome devaluation.

#### Secondary parallel (whole-block) fMRI analysis

We repeated the same task-fMRI analyses while modelling the entire blocks (rather than their onsets) at the first-level GLM. To account for multiple comparisons, we used a stricter cluster significance threshold of p=0.001. Running the within-group analyses in the extensive training group yielded a significant correlation between training duration-induced changes in neural activity (as modelled using a linear trend analysis) and the behavioral habit index in the right cuneal cortex (cluster size=147, max Z-value =4.20, cluster-corrected p<0.001; Supplementary Figure 4). This effect was driven by greater activations in this region during rest compared with task blocks in early stages of the task (and became indifferent in later stages) in habitual participants whereas in goal-directed participants it was indifferent throughout the entire task.

In addition, when testing the correlation between the behavioral habit index and the change in BOLD response toward still-valued vs. devalued stimuli (throughout the entire blocks) following outcome devaluation we found an abundance of clusters in motor, somatosensory and visual regions (see all identified clusters in Supplementary Table 2). The more participants demonstrated goal-directed behavior, the larger the difference in the response in these regions was. This was not surprising as it putatively reflects the fact that goal-directed participants are those that had reduced their relative responding toward the devalued vs. the valued outcomes.

### Subgroup-based exploratory analyses

In our pre-registration we considered the possibility of null results and stipulated that in such a case we will cluster participants within the two training groups into habitual and goal-directed subgroups to examine related neural differences. As noted above, when observing the behavioral data from the pre-registered analyses we noticed that the devaluation-induced effects indicating goal-directed behavior in both groups were driven only by a distinct subset of participants, whereas many participants in both groups actually responded habitually. This further motivated the following goal-directed vs. habitual subgroup analysis approach.

#### Clustering participants into subgroups based on their habit index

In this exploratory analysis we performed the following steps: (1) We removed extremely unengaged participants (with < 3SD response rate during the last training run from their group average). This resulted in the removal of three participants from the extensive training group. (2) A decrease in responding towards both outcomes to some extent could be probably best explained by a general motivational effect and thus be aligned with the interpretation of habitual behavior. However, our habit index calculation would also characterize participants who extremely reduced their response rate toward both outcomes as habitual. We surmise such cases most likely represent goal-directed behavior that accounts for general satiety. Thus, we re-calculated the habit index for participants who reduced their responding towards both snacks in more than 90% following outcome devaluation, by measuring the difference between the mean response rate toward both snacks after devaluation minus the same measure before devaluation. (3) Following previous work by some of the coauthors of this manuscript^87^, to identify latent subgroup in each training group we ran a finite mixture model on the habit index using the Flexmix R package^88^. We ran the model using k=1 to 5 clusters and found that two clusters within each group were best suited to explain the habit index data (based on the lowest Bayesian Information Criterion; Figure 8). The two clusters appear to separately capture habitual participants (with values around 0) and goal-directed participants (with larger values). (4) As a final step we exploited the results of the cluster analysis to identify and remove a minority of participants who appeared to respond irrationally (responding much more for the devalued compared to the still-valued outcome during the extinction test). To achieve this, we removed participants that were classified in the “goal-directed” cluster despite their habit index being lower (rather than higher) than participants in the “habitual cluster” (See the left tail of the “goal-directed” clusters in Figure 8). This resulted in the removal of three participants.

**Figure 8.**
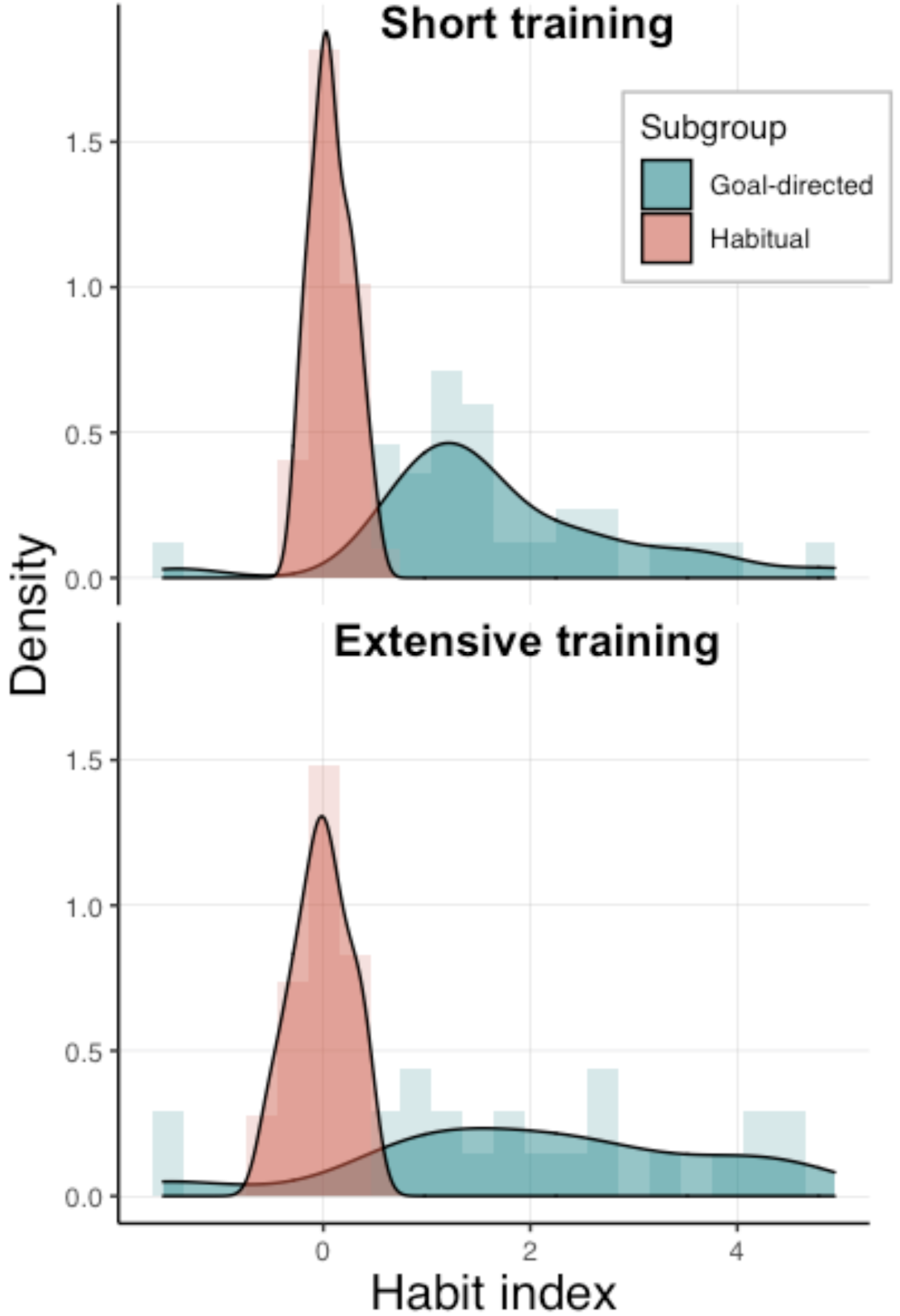
Cluster analysis. The behavioral habit index is best explained by two latent sub-groups identified within each group, as shown by a finite-mixture modelling analysis (testing k=1 to 5 potential clusters and comparing their Bayesian Information Criterion). The sub-groups distinguish habitual participants (with scores around 0) from goaldirected participants (with larger scores).

Following this process, we now had 57 participants in the extensive training group (21 goal-directed and 36 habitual) and 60 in the short training group (27 goal-directed and 33 habitual). This classification was then used along with the neuroimaging data to compare these inferred subgroups within each training condition to identify related functional and micro-structural differences.

### Subgroup-based task-fMRI analysis

#### Training duration-induced functional plasticity and tendency to form/express habits

We compared our inferred goal-directed and habitual subgroups within the extensive training group on each of the training duration-related contrasts (see examined contrasts 2-5 in Table 1). Whole-brain analyses revealed larger/increased training-induced BOLD activations in habitual participants relative to goal-directed participants in the bilateral intra-parietal sulcus and surrounding regions. These clusters were identified both when comparing the last two vs. first two training runs (left: cluster size=136, max Z-value=4.29, cluster-corrected p<0.001; right: cluster size=92, max Z-value=4.53, cluster-corrected p<0.001), as well as when comparing linear trends across the entire task (left: cluster size=211, max Z-value=4.01, cluster-corrected p<0.001; right: cluster size=94, max Z-value=4.27, cluster-corrected p<0.001; Figure 9A), suggesting this region is implicated in the tendency to form habits. In the latter analysis we also identified a third adjacent cluster in the left supramarginal gyrus (cluster size=77, max Z-value=4.02, cluster-corrected p=0.004), and interestingly, a cluster in the dorsolateral-prefrontal cortex (dlPFC; cluster size=59, max Z-value=4.65, cluster-corrected p=0.017). These effects were mainly driven by a decrease in activation in goal-directed participants from initial to later stages of training (see Figure 9B for example) suggesting the existence of early differences in neural processing between participants who were more likely to from and express habits vs. those who tended to respond goal-directedly or were able to overcome habit expression when it was not adaptive. Interestingly, we did not find any effects in the opposite direction, that is, regions that relatively increased their cue-reactivity in the goal-directed subgroup compared with the habitual subgroup. Additionally, analyzing the same contrasts using SVC analysis of any of our ROIs did not yield a significant effect. Finally, note that in contrast to our individual-differences analysis, we did not find any differential within-day effects between the two subgroups.

**Figure 9.**
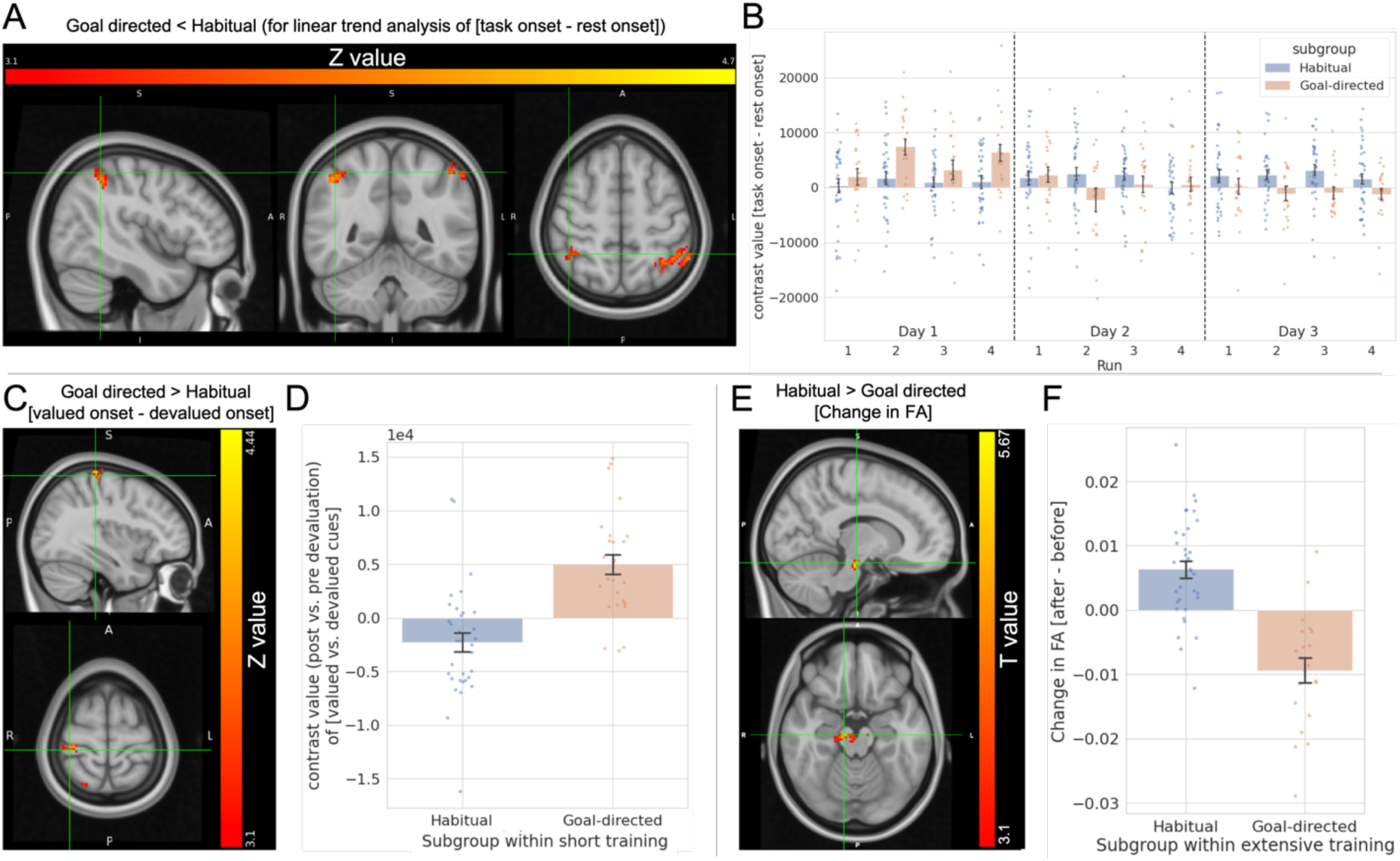
Exploratory analysis of differential functional and microstructural plasticity in inferred subgroups of habitual and goal-directed participants. (A) Whole-brain analysis testing linear trends in activation across training of [task blocks onset vs. rest blocks onset] between habitual and goal-directed subgroups within the extensive training group. A linear trend was modelled by assigning linear weights to the training runs. The bilateral intraparietal sulcus reduced its activation in goal-directed participants compared with habitual participants as training progressed. (B) Plots are presented for illustrative purposes of the effects in panel A: averaged contrast values of task onset vs. rest onset within the identified significant clusters in the bilateral intraparietal sulcus (shown in panel A) in both subgroups. (C) Whole-brain analysis comparing changes in BOLD response toward the still-valued vs. devalued cues following outcome devaluation between habitual and goal-directed subgroups within the short training group. The right postcentral gyrus increased its activity toward the still-valued snack in goal-directed participants compared with habitual participants. (D) Plots are presented for illustrative purposes of the effects in panel C: averaged contrast values of the change of [valued vs. devalued block onsets] following outcome devaluation (after vs. before) in two significant clusters identified in the right postcentral gyrus (one of them is shown in panel C). (E) Whole-brain analysis comparing training-induced changes in FA between habitual and goal-directed subgroups within the extensive training group. We identified a cluster within the ventral tegmental area (VTA) and substantia nigra (SN) in which FA increased in habitual and decreased in goal-directed participants. (F) Plots are presented for illustrative purposes of the effects in panel E: averaged contrast values within the significant FA cluster in the VTA and SN (shown in panel E). Notes: (1) All DTI and fMRI results were cluster-corrected (p<0.05). (2) The subgroups partition was based on finite mixture modelling of the individual habit index calculated for each participant based on their task performance. (3) Error bars in bar plots indicate 68% confidence interval (equivalent to ±1 std error).

#### Neural correlates of differential value-based cue reactivity and tendency to form/express habits

We compared changes in neural reactivity to valued vs. devalued cues following outcome devaluation between the two subgroups (separately within each of the training groups). While we did not find any difference in the extensive training group, we have identified 3 clusters in the short training group indicating increased activation for the valued vs. devalued snacks following outcome devaluation in the goal-directed subgroup compared with the habitual subgroup (right superior parietal lobule: cluster size=80, max Z-value=3.8, cluster-corrected p<0.001; right postcentral gyrus cortex: cluster size=72, max Z-value=4.38, cluster-corrected p=0.002 and cluster size=46, max Z-value=4.4, cluster-corrected p=0.027; Figure 9C). These results imply that a tendency to quickly acquire and express habits is related to reduced neural sensitivity to value-related changes (Figure 9D).

### Subgroup-based DTI analysis

#### Training duration-induced micro-structural neural plasticity and tendency to form/express habits

We compared the training-induced changes in MD and FA between habitual and goal-directed subgroups within each experimental group. Whole-brain analysis in the extensive training group revealed different patterns of training-induced changes in FA in the ventral tegmental area (VTA) and substantia nigra (SN) between the habitual and goal-directed participants (cluster size=249, max t_53_=5.62, cluster-corrected p=0.031; Figure 9E). This effect was driven by an increase in FA in habitual participants and a decrease in goal-directed participants (Figure 9F), suggesting that differential micro-structural changes in midbrain dopaminergic regions may tilt action control to be dominated by one system or the other. We did not identify any differential micro-structural changes in the short training group.

### A note on further exploratory analyses

The scope of this work is comprehensive and included multi-modal MRI data analysis (task-fMRI and DTI) at several levels (individual, within- and between-group analyses). Additionally, the pattern of our results led us to perform a substantial exploratory analysis in addition to other planned secondary analyses. Therefore, for the sake of focus and clarity we have decided not to include in the scope of this work exploratory analyses of the resting-state fMRI data and of the two-step sequential decision-making task.

## Discussion

In this registered report, we first set out to examine the capacity of the free-operant habit induction paradigm used by Tricomi et al.^23^ to induce habits in humans as a function of training duration and whether the putamen increases its cue reactivity accordingly, as was found in the original study. Similar to previous attempts to induce habits using this procedure in behavioral settings^23,87^, we did not find the hypothesized effect of decreased devaluation sensitivity (as an index of habit formation) following extensive compared with short training. Surprisingly, we found that in the original contrast used, the putamen reduced, rather than increased, its activity following extensive training to cues associated with a rewarded action. Nevertheless, focusing specifically on the posterior putamen (in an exploratory follow-up analysis), the sub-region of the putamen in which Tricomi et al. ^23^ found the increased activation (the posterior part), we found an effect trending in the same direction as in the original study (but given the absence of the main hypothesized behavioral effect we cannot relate it to habits). Furthermore, we used our relatively large sample, to identify functional and micro-structural neural correlates of individual habit expression. We found that within-day increased activity (that is, an increase across each of the three training days throughout the daily training, averaged across days) in the left head of caudate toward cues paired with rewarded actions was associated with goal-directed behavior and vice versa. Additionally, we found that a low-level visual region (the left intracalcarine sulcus) was more reactive toward devalued vs. still-valued cues in goal-directed participants but not in habitual participants. We did not identify any micro-structural plasticity (as indexed by changes in MD and FA) related with individual habit expression. Finally, as planned for the case of null behavioral effect, we conducted an exploratory analysis comparing goal-directed and habitual subgroups (clustered based on the behavioral data) within each experimental group. We found a reduction in activation in the bilateral intraparietal sulcus towards cues associated with rewarded actions as training progressed in goal-directed participants but not in habitual ones (within the extensive training group). Additionally, within the short training group, we found that parietal regions (right superior parietal lobule and postcentral gyrus) became more reactive toward still-valued vs. devalued cues following outcome devaluation in goal-directed participants compared with habitual participants. Finally, we found differential extensive training-induced changes in FA between habitual and goal-directed participants in midbrain dopaminergic regions (VTA and SN), such that FA levels increased in habitual participants and decreased in goal-directed ones.

### Tricomi et al.’s (2009) paradigm and experimental habit induction in humans

The results of our pre-registered behavioral analysis are consistent with the results of the two replication attempts of Tricomi et al.’s^23^ behavioral findings, conducted by de Wit et al.^38^ (two attempts) and a recent multi-laboratory work by Pool et al.^87^. In the latter, the researchers conducted a large-scale replication attempt of this paradigm using five different samples across four different sites. In both de Wit et al.^38^, Pool et al.^87^ and the present work, the statistical analysis demonstrated that participants in both groups were sensitive to outcome devaluation, manifested as a relatively greater reduction in responding toward the devalued compared with the still-valued outcome (although this effect was driven by a small subset of participants whereas most participants were insensitive to outcome devaluation; see more below). Importantly, in contrast to the original Tricomi et al. findings^23^, participants’ sensitivity to devaluation was not reduced following extensive training. Note that Pool et al.^87^ found that the affective component of stress moderated the training duration effects on sensitivity to outcome devaluation. Specifically, participants low on this measure demonstrated reduced sensitivity to devaluation following extensive vs. short training, whereas participants high on this measure responded habitually already after a short training. In an exploratory analysis, we examined if this effect was present in our data as well. Although not conclusive, we found a trend indicating a moderating role for the affective component of stress on habit formation as a function of training extension (for an elaborated discussion on this potential moderating role see ^87^). As Pool et al. had a substantially larger sample, the fact that this effect was only trending in the present data may reflect reduced power of this smaller sample size. Still, taken together, the present work together with the aforementioned previous replication attempts form a solid body of evidence suggesting that the Tricomi et al. paradigm^23^ is of limited utility in terms of its capacity to elicit training-related effects on the behavioral expression of habits in a robust manner.

Particular contributions of the present work to determining the efficacy of Tricomi et al.’s procedure^23^ include the involvement of two of the original study’s coauthors (first and last authors) which helps to prevent significant misalignments of the methods and analyses between the original and the present work^89^. Another important and unique contribution of this work is that, while the previous replication attempts were behavior-only studies, in the present work the behavioral task was performed during fMRI scans, similar to the original study. As mentioned above, Pool et al.^87^ found that participants with lower rates of the affective component of stress (which includes anxiety) exemplify the hypothesized effect of training extension on habit formation. In light of recent evidence showing lower rates of anxiety in participants enrolling in fMRI studies compared to participants enrolling in behavioral studies^90^, it has been suggested that a self-selection bias may have been responsible for the contradictory results obtained when utilizing the Tricomi et al.’s procedure^23^ in different experimental settings^87^. Specifically, in the behavior-only replication attempts participants may have been self-selected to have higher anxiety rates compared with participants in the original fMRI study. Such an explanation could have potentially reconciled the discrepancies between the effect observed in Tricomi et al.^23^ and the lack of similar effects in De Wit et al.^38^ and Pool et al.^87^. Here, by using an fMRI setting, we tested this possibility directly and our results provide evidence that the experimental setting along with potential anxiety-based self-selection biases are not sufficient to account for the discrepancy between the original study and subsequent replication attempts. Whether or not a selection bias does occur in a given fMRI study will depend on the specific recruitment procedures used at particular research centers, and thus it remains possible that this bias did contribute to differences between the behavioral effects observed in the current and the original study. However, it is most parsimonious to assume that either the main behavioral result of the original study was a false positive or that the discrepancies in results are attributable to other unknown factors.

Similar to Pool et al.^87^, and largely consistent with the results obtained by De Wit et al. when the researchers used the same procedure^38^, close inspection of the data obtained in our study (supported by empirical evidence obtained by the cluster analysis of the habit index) revealed an interesting finding. In both groups, while the statistical analysis indicated the presence of sensitivity to outcome devaluation and thereby the dominance of goal-directed behavior, in practice, only a small subset of participants that responded in a goal-directed manner drove this effect, whereas the majority of participants responded habitually (i.e., were insensitive to outcome devaluation). This suggests, perhaps counterintuitively, that this paradigm fails to exemplify habit formation as a function of training duration due to it being too prone to rapidly induce habits, instead of not being able to induce habits following extensive training. At least as it is currently being used, this paradigm appears not to be sensitive enough to capture sufficient variability in sensitivity to outcome devaluation. Participants’ rapid development of habit-like behavior (insensitivity to outcome devaluation) hinders the detection of effects induced by the additional training sessions in the 3-day group. Therefore, future efforts to change this procedure or establish a new one to reliably induce habits through training extension, may benefit from initially focusing on methods to prevent rapid manifestation of insensitivity to outcome devolution. One clear target for modification in future studies is the reinforcement schedule. The Tricomi et al. paradigm^23^ uses a variable interval (VI) reinforcement schedule which has been shown to enhance habit formation compared with variable ratio (VR) reinforcement schedule in rodents^91–94^. While the capacity of these two reinforcement schedules in forming habits have not been directly compared in humans, our evidence may imply that a similar effect in humans exists and motivates the use of VR schedules in future studies in order to avoid rapid habit formation. Furthermore, based on other studies and the present study, it appears that habit formation and expression vary considerably across individuals and seem to be susceptible to multiple individual differences factors and moderating effects (e.g., ^24,87,95^). Investing more effort in identifying these factors and, in turn, accounting for them, may be necessary in order to establish experimental habit induction in humans.

Another possible limitation of this paradigm^23^ is that it is unclear whether the devaluation test as implemented, is sufficient to adequately distinguish goal-directed from habitual behavior. Using a test that can confidently attribute “goal-directedness” to observed actions (to ensure that an unchanged response pattern following outcome devaluation is indeed habitual) was recently suggested as a criterion that has to be satisfied in order to reliably interpret the results of research in the field^96^. It is possible that the procedure we used here does not satisfy this criterion for various possible reasons, including the following: (1) Participants were most likely aware that they did not have to eat their potential devalued winnings during the test session; (2) Responding during the test toward the devalued outcome imposed little to no cost at all on participants. The only cost was the supposedly physical effort of pressing a keyboard button. We suggest that it is likely that at least some of the participants may have responded toward the devalued outcome simply as it had no tangible cost and the only viable alternative was to sit and stare at a static fractal image (which may not be preferable for some); (3) Similarly, the potential cognitive effort of withholding the signaled response may have been larger than the physical effort of pressing the button, and these two types of efforts may have been thoughtfully considered in a goal-directed manner rather than being automatically executed. Therefore, we suggest that future attempts to establish experimental habit induction procedures in humans could put more of an emphasis on aligning participants’ incentives with their actions. To achieve this, future procedures could incorporate a tangible cost imposed on each action (see^95^ for such implementation in the context of the two-step task with incorporated outcome devaluation) and allow the utilization of other actions beside choosing between responding or doing nothing.

### The putamen’s role in training and habitual action control

A key neural hub thought to be dominant in acquiring and executing habitual behavior is the putamen^3^. This motivated us to place it in the center of various analyses conducted as part of this research and to hypothesize it would be correlated with habit expression at both the group and individual levels. Comparing cue-related BOLD activations in early vs. final stages of extensive training, similar to the analysis conducted by Tricomi et al.^23^, did not reveal a similar effect in the putamen. This, by itself, can be simply explained as consistent with the lack of stronger habit formation following extensive training observed in the current work (in contrast to the Tricomi et al.’s finding^23^). However, not only did we not find an increase, but the bilateral putamen reduced its BOLD activation as training progressed. Notably, while this result is consistent with a body of research reporting a reduction in the putamen activation as training progresses (e.g., ^97,98^, for review see ^99^) it is inconsistent with Tricomi et al. and other research showing increased activity in the putamen as training progresses^23,100–102^. This discrepancy can be reconciled by taking into consideration the specific subregions in the putamen where the activations were observed. In the Tricomi et al. study^23^, the posterior putamen was specifically found to show an increasing activation profile with training. This region is anatomically distinct from the more anterior aspects of the putamen in which the decreasing activation profile was found in the present study. In an exploratory analysis focusing on the smaller specific region of the posterior putamen reported by Tricomi et al.^23^, we did find a statistical trend toward increasing activity as a function of training (albeit not quite reaching statistical significance). Accordingly, differential response profiles between the anterior and posterior parts of the putamen were also clearly observed when visually inspecting raw training-induced activations across the bilateral putamen. Those results could be interpreted as being broadly consistent with previous literature finding decreasing activation in the anterior (associative) putamen and increasing activation in the posterior (sensorimotor) putamen with training^23,97,98,100–102^. Indeed, a transfer of motor representations is suggested to occur from anterior to posterior putamen as a function of training^103–105^. These opposing effects have not often been seen side by side in the same study and further research is needed to disentangle the specific role of different putamen subregions and their time-dependent engagement patterns in human action control. Perhaps a further functional subdivision of both the associative and particularly the sensorimotor putamen is required to gain a detailed understanding of the putamen’s contribution to instrumental learning. In the present study, it should be emphasized that due to the lack of behavioral effects of training duration on sensitivity to outcome devaluation, any training duration-induced changes in the putamen observed at the group level could not be attributed to the expression of habitual/goal-directed behavior but solely to the effects of training.

Contrary to our hypotheses, we did not find any evidence for functional or microstructural plasticity in the putamen associated with individual habit expression (measured as sensitivity to outcome devaluation), not even when we clustered participants into goal-directed and habitual sub-groups. In rodents, convincing evidence dissociating the contributions of the dorsomedial striatum (homologous to the human anterior caudate) to goal-directed behavior and dorsolateral striatum (homologous to the human putamen) to habitual responding^28,32^ have spurred the idea of a shift in action control between these regions (from the former to the latter) as goal-directed behavior shifts into habits^106^. Similarly, it is thought that a corresponding shift from goal-directed to habitual behavior in humans predominantly results from a transition in action control from the anterior caudate to the putamen^3^. While some evidence, although correlational, supports this idea^23,37,107^, our findings join a growing body of findings that do not find evidence for such a transition in healthy individuals^98,108–110^. Notably, targeting individual differences also yielded mixed results regarding the putamen’s role in determining habitual performance. While De wit et al.^36^ found that putamen-premotor cortex white matter tract strength, as well as putamen gray matter density, were associated with habitual slip of actions in healthy individuals, other more recent studies have not found functional or structural neural correlates between the putamen and individual differences in habit expression^109–111^. One potential explanation could be that the procedures used to operationalize and measure habits do not capture well or in a consistent manner the construct they intend to. This could lead to mixed results regarding the neural substrates of human habitual control. Alternatively, habit expression may be subject to multiple environmental and psychological transient as well as stable factors, rendering related effects subtle and unstable. In that eventuality, even if a more effective task paradigm were to be used, large sample sizes are likely to be required to reliably implicate and characterize the underlying neural mechanisms. Relatedly, the striatum may be differentially recruited as a function of the nature of the task involved^112^. Tasks employing a sequence of actions often find increased activation in the sensorimotor striatum (e.g., ^97,103^), whereas single-motor action tasks in rodents report a training duration-related reduction in the amount of task-related active neurons in this region (e.g., ^113,114^). Helie et al.^99^ suggested that increased activation in the sensorimotor striatum following extensive training occurs after response selection and is in fact associated with response-response associations required for automatically executing a sequence of actions. Outcome devaluation studies in humans typically use a single action, such as a simple button press, which may thereby not be controlled by the putamen or alternatively, introduce a very subtle effect, frequently undetected. Finally, we cannot rule out the possibility that the role of the human (posterior) putamen in habitual control is not equivalent to the role of the dorsolateral striatum in rodents as commonly assumed. It is possible that humans’ advanced ability to engage in goal-directed planning and execution implies and/or generates different neural implementations of habitual control. Alternatively, changes in neural regions and circuits associated with goal-directed behavior as behavior becomes habitual may be more robust and thus more readily captured in the current available methods, as seen in the present and other studies (e.g., ^111^). As a final note, it is certainly possible that the putamen plays a role in habit learning but that whether or not habits are expressed in a given situation is contingent on factors or gated by brain regions that are not directly associated with the level of training-induced functional or microstructural plasticity in the putamen.

### Neural correlates of the individual tendency to form/express habits

We found that patterns of daily activation changes in the left head of caudate (but not changes across the entire three days of training) predicted individuals’ sensitivity to outcome devaluation. This is consistent with the notion that sensitivity to outcome devaluation may be manifested in differential neural activity in goal-directed regions (e.g., ^111^). This finding is also aligned with evidence relating caudate-vmPFC white matter tract strength^36^ and caudate increased activation during trials involving a valued outcome^109^ with individual goal-directed action control. An important distinction regarding the novelty of this finding is that it illustrates the importance of re-occurring patterns of within-session activation changes in general and the implication (through such a process) of the head of caudate in maintaining goal-directed behavior. Our results suggest that each training session is to some extent a new context that re-initiates the engagement of the head of caudate. Decreased reactivity of this region is indicative of reduced sensitivity to outcome devaluation whereas an increase is associated with goal-directed behavior. This finding perhaps implies that if the training would have continued (i.e., involve further sessions), at some point (varied across individuals), as the slightly new contexts that each day constitute generalize (and a habit is simultaneously formed), the activation patterns in the head of caudate of the initially more “goal-directed” participants would become similar to those of the “habitual” participants and render their behavior habitual. This idea is supported by research showing that the caudate encodes the contingency between actions and outcome^115,116^ which is crucial for goal-directed behavior. However, our results here are correlational and this idea is speculative and should be directly tested to be verified. Yet, if indeed verified, the head of caudate may putatively be used as an online within-session/day marker of habit formation. Notably, in contrast to the caudate, we did not find any functional or structural effects associated with sensitivity to outcome devaluation in the vmPFC. This region has been largely implicated in goal-directed behavior (e.g., ^4,35–37,117^). Perhaps as most participants in our study had become habitual already after a short training, there was not enough variance in the data to observe an effect in the vmPFC. This region is known to have a major role in processing the value of an outcome (e.g., ^118^), which, taken together with the effect found in the head of caudate, may indicate that in the current procedure value signals are less predictive of action control than contingency signals.

Another novel finding of this work is that individual sensitivity to outcome devaluation involves differential processing of valued vs. devalued cues in the left primary visual cortex. Goal-directed behavior was associated with greater activation in this region toward a devalued cue (as a function of the value reduction). This suggests that action control is directly related to differential neural processing of predictive stimuli in lower-level sensory regions and thus during earlier stages of the neural processing of external information than had been previously thought. To the best of our knowledge, to date only one study implicated visual regions in goal-directed/habitual behavior in humans^25^. That research demonstrated greater activation in the middle and superior occipital cortex toward a condition that involved stimulus-response encoding (which presumably underlies habitual action control) vs. a condition that encouraged the utilization of response-outcome associations (which presumably underlie goal-directed action control). However, that study implicated different occipital regions (of higher visual processing levels) than the regions identified in the current study (the primary visual cortex). Also, as there are known projections from the primary visual cortex to the dorsomedial striatum^119^ (the rodent homologue of the human caudate), it is possible that the differential processing of cue values in this region contributes to the effect we found in the left head of caudate. Finally, this finding can be possibly integrated with the exploratory finding showing that (within the short training group) the right postcentral gyrus and right superior parietal lobule were more reactive to valued vs. devalued cues in goal-directed participants but not in habitual ones. In primates, the superior parietal lobule is suggested to integrate visual and somatosensory information to control goal-directed actions executed by the arms (See^120^ for review). Taken together, it is possible that increased sensitivity to devalued cues in low-level visual regions in individuals with higher tendency for goal-directed behavior is, at least to some extent, directly associated with the increased sensitivity in the postcentral gyrus and superior parietal lobule. For example, the increased activity toward devalued cues in the lower-level visual regions may inhibit activations in the superior parietal lobule and postcentral gyrus, which in turn results in a differential activation in favor of the valued cues in these regions.

Our subgroup-based analysis, in which we divided participants in each experimental group to goal-directed and habitual subgroups based on their demonstrated sensitivity to outcome devaluation, has related a few other regions of both the sensorimotor and associative parts of the cortico-striatal circuits^106,121^ to habit expression. We found that the bilateral intraparietal sulci, the left supramarginal gyrus (part of the inferior parietal lobule) and the dlPFC decreased their activity as training progressed in participants that maintained goal-directed behavior (but not in those that responded habitually) following extensive training. This decrease in activation was derived from elevated activation levels in early stages of training in goal-directed participants but it was absent in habitual participants. Both the inferior posterior parietal lobe and dlPFC play a major role in goal-directed behavior by implementing related required processes. They both constitute a prominent part of the fronto-parietal brain network (a component of the cognitive control network^122^) which facilitates cognitive control and flexible adaptation to task demands (adaptive control)^123^. They have also been directly associated with planning processes^102^, have shown to be involved in state space representation (and model-based learning)^124^ and were implicated in stimulus-dependent attentional shift (exogenous adjustment)^125^.

The role of these regions (the inferior posterior parietal lobe and dlPFC) in the context of habit learning was highlighted by a few studies. Evidence implicating the inferior parietal lobule in sensitivity to outcome devaluation was provided by Liljeholm et al.^25^ that found that greater discriminatory activation in this region between stimulus-response and response-outcome pairings predicted sensitivity to outcome devaluation. Contradictory findings (at first glance) to our work were found by Poldrack et al.^97^ and Zwosta et al.^111^. Poldrack et al.^97^ found practice-related decreased activity in the both parietal and dlPFC regions, which were indirectly associated with the formation of behavioral automaticity (as indicated by dual-task interference elimination). Nevertheless, that study focused on skill learning and used the serial reaction time (SRT) task^126^ (either in single or dual-task conditions) which demands substantial cognitive resources, especially when compared to our non-challenging free-operant task. It is possible that when high cognitive resources are required, the dlPFC and inferior parietal lobules are more engaged in early learning stages also in individuals who tend to quickly express automatic behavior. Another key difference is that automaticity in the SRT task is constantly desired, that is, participants are not placed in a condition aimed to examine their capacity to adjust their learned behavior (in which they benefit from overcoming acquired automaticity). Goal-directed participants in our procedure may have differentiated from habitual participants not by their capacity to engage in habitual action control but rather by their ability to switch back when the circumstances have changed.

Zwosta et al.^111^ found that stronger habit expression was associated with a larger decrease in the inferior parietal lobule across training, an effect in the opposite direction than reported here. Notably, that task’s operationalization of the targeted neurobehavioral processes was very different from the one we used. It is composed of binary choice trials and change in explicit goals between task phases. Thus, the possibility of substantial involvement of the inferior parietal lobule also in participants that tended to quickly express habits, apply in this/that work as well. Importantly, that work employed a relatively short training (across one day) and therefore focused on changes within a single session rather than across multiple (spaced) sessions. This raises the possibility that across different timescales different patterns of functional plasticity in the inferior parietal lobe play a role in habit learning. Another important emphasis regarding the work by Zwosta et al.^111^ is that habitual behavior was not indexed as sensitivity to outcome devaluation but rather as increased error rates and increased response time in trials that were incongruent with respect to previous learning strategies (contrasting putative goal-directed and habitual action control). The effect in the inferior parietal lobule was only observed for the response time measure but not for the error measure. Notably, the latter is more equivalent to responses that are insensitive to outcome devaluation than the former.

Our work provides novel evidence that significant recruitment of the inferior parietal and dlPFC regions at early stages of training can lead to larger flexibility to adjust instrumental behavior when necessary in later stages. Thus, while Zwosta et al.^111^ speculated that individual differences in habit expression are mediated by inferior parietal lobule activations, which reflects the anticipation for a specific outcome at the end of training, our results suggest that there might be a critical phase at the beginning of training in which activations in this region may take part in forming the neural machinery required for later behavioral adaptations. The later use of this machinery appears to no longer require the particular involvement of the inferior parietal lobule. Such interpretation implies that activation in these regions can perhaps be used as an early-stage neural marker of the likelihood of recruiting goal-directed action control. Such neural marker could putatively be used to identify individuals’ tendency to form and express habits. More research is needed to understand whether such activations are associated with a specific state or are stable within individuals, namely reoccur over similar and different tasks, and thus represent an individual trait.

The cognitive and mental processes relating these early activations with later goal-directed performance can also be a target for future exploration. A possible explanation for the role of these early-stage activations may rely on other related findings about the role of these regions. Sohn et al.^125^ implicated several parietal regions, including the posterior inferior parietal lobe in task switching. Glascher et al.^124^ have shown that activity in the dlPFC reflects state prediction errors, thereby having a major role in encoding the model of the environment in model-based reinforcement learning, and Mcnamee et al.^37^ found that this region encoded information about both the identities of actions and outcomes, suggesting it is involved in encoding such model. Taken together, it may be that increased activations in the dlPFC and inferior parietal sulcus generate a good representation (model) of the task states, action and related outcomes, and perhaps generate a different “sub task” representation for each associative structure. Establishing and rehearsing these representations at initial stages of training may have made it easier in later stages to distinguish between the different conditions (valued vs. devalued) once it had become beneficial to do so. Another important observation, made by Zwosta et al.^111^ and reinforced by the current study is that individual differences in habit expression appear to mainly stem from individual differences in regions associated with goal-directed rather than habitual action control. This may indicate that individual differences in regions of the goal-directed system are perhaps more common and more substantial than in regions of the habitual system or alternatively, that their effects on behavior are stronger.

Another novel finding of this work is the identification of differential extensive training-induced microstructural changes in midbrain dopaminergic regions (VTA/SN) and their surroundings between goal-directed and habitual participants. FA was found to elevate in these regions for habitual participants and was reduced for goal-directed ones. Dopamine is thought to play a crucial role in habit formation and in moderating the balance between goal-directed and habitual action control^3,127^, but the exact mechanisms by which it acts are not yet fully understood. Most notably, research in animals and humans often ascribe a different, even opposing, role for dopamine in action control. Whereas dopamine has been shown to promote habit formation in rodents^128,129^ (but note the D1 and D2 receptors’ dissociable effects, promoting and depressing habit formation, respectively^130^), it was found to enhance goal-directed behavior (and model-based learning)^131–135^ in humans (but see ^136^). As far as we know, habit research in humans has yet to uncover direct associations between habit formation and structural or functional plasticity in dopaminergic regions but rather only in their efferents’ target regions. We show here that different patterns of microstructural plasticity in midbrain dopaminergic regions may underlie shifts in action control or the ability to adjust behavior when the goal of a behavior ceases to be desired. It is important to note here, that since (1) this region is comprised of both white and gray matter and the cluster found covers a relatively large area, (2) we did not find MD changes and (3) the FA measure can be affected by various factors in the tissue microstructure^137^, any attempt to interpret the underlying neural and cognitive processes associated with these microstructural changes would be highly speculative and further focused research is required to characterize these processes. Based on our findings, the only suggestion we can make in this context is that individual tendency to acquire and express habits may stem from differential dopaminergic modulation of goal-directed and habit neural circuits (involving the associative and sensorimotor striatum, respectively). As fMRI BOLD activations are mainly a product of neural processing localized around the afferent inputs region^138^, in this work, by using DTI indices, we might have traced back the source of activations in cortical and striatal regions identified as implicated in goal-directed action control that dissociates individuals with different habit learning tendencies. It is important to emphasize with respect to all of the findings obtained through the subgroup analysis that this analysis, although conceptually planned for the case of null behavioral effect, is exploratory and as such its results should be further validated in targeted future research.

### Conclusions

In conclusion, in this well-powered registered report, we provide further evidence that the Tricomi et al. paradigm^23^ is not well-suited to reliably demonstrate the effect of habit formation as a function of training duration, potentially due to a rapid manifestation of habit-like behavior, which is expressed already after short training. Furthermore, we were not able to relate any functional or microstructural plasticity in the putamen with individual habit expression. Instead, regions commonly implicated in goal-directed regions were most predictive of individual habit expression. We show that re-occurring within-day increased activations in the head of caudate throughout training and early-stage elevated activations in frontoparietal regions (mainly in the inferior parietal lobe and the dlPFC) were associated with goal directed behavior (and vice versa). Additionally, we found evidence that larger neural reactivity toward devalued cues in the primary visual cortex and increased reactivity in favor of still-valued cues in somatosensory and superior parietal regions are related to individual tendencies to express goal-directed behavior (and vice versa). Finally, we found that differential patterns of training-related microstructural plasticity in midbrain dopaminergic regions were indicative of habit expression. Taken together, this work provides new insights regarding the underlying putative neural mechanisms involved in individual habit learning and motivates the development of novel, well-informed, procedures for experimental habit induction in humans in order to accelerate research in the field.

## Declarations of interest

None.

## Acknowledgments

We are thankful to Jasmine Segal, Miri Goldman and Danielle Cohen who helped us with recruitment and data collection.

## Funding

This work was supported by the European Research Council (ERC) [grant number 715016]; and the Israeli Science Foundation [grant number 1798/15 and 1996/20]. Rani Gera was supported by the Fields-Rayant Minducate Learning Innovation Research Center.

## Notes

### Competing Interest Statement

The authors have declared no competing interest.

