## Supplementary materials for "Characterizing habit learning in the human brain at the individual and group levels: a multi-modal MRI study"

#### Extension and elaboration on the exploratory examination of stress affect's moderating role of the effects of training duration on habit formation

*Exploratory factorial analysis*

As we did not find a training duration effect on habit formation and since, similar to previous research[1,2], there were evidence that the majority of participants have demonstrated insensitivity to outcome devaluation even after a short training, we followed the recent work by Pool et al.[1] and set to explore whether the affective component of stress moderates the effect of training extension on habit formation. Out of the questionnaire data we collected, to adhere with Pool et al.[1] we used the BIS-11, TICS and STAI questionnaire in an exploratory factor analysis (EFA) aimed to identify significant factors that underlie these questionnaires' data. Note that in Pool et al.[1] each participant filled out either the SATI-state or the STAI-trait questionnaire (varied across samples) and their data was used interchangeably, here we collected data for both questionnaire and thus we averaged their score for each participant. The TICS included nine subscales and the BIS-11 included three. Thus, respectively, combined with the STAI score, each participant had 13 questionnaire scores entered into the EFA. For the EFA we used oblimin rotation as implemented by the R "psych" package[3].

Running the "parallel" methos implemented by the "psych" packzge[3] suggested that three underline the data. Note that this is different from Pool et al.[1] where this analysis suggested four factors, a difference that may stem from the difference in sample sizes (n=199 in Pool et al.'s analysis vs. n=123 in the present work) . We then run the EFA and labeled as "stress affect" one of the factors, which largely resembled the equivalent factor identified by Pool et al.[1] (see Supplementary Table 1).

*Testing the role of stress affect in habit formation as a function of training duration*

* The following part is adapted from the main text and is added here for fluency and completeness.

We included the factor most corresponding with stress affect along with Devaluation (valued or devalued outcome), Group (1-day or 3-day) and all possible interactions as independent variables in a mixed-effects linear regression model where the dependent measure is the change in response rate following outcome devaluation. We used participant as a random factor. We found a marginally significant interaction between the stress affect, devaluation and group ($\chi^2_1$ = 3.55, p=0.059), indicating stress affect may modulate the interaction between devaluation and training length. We followed this analysis with a simple slope approach and tested the interaction between devaluation and group for participants low (-1 SD) and high (+1 SD) on the stress affect measure. We found a marginally significant effect for participants with low levels of this measure ($\beta$= -0.38, 95%CI [-0.82, 0.06], $p$=0.097) and no significant effect of those with high levels ($\beta$= 0.24, 95%CI [-0.21, 0.68], $p$=0.304; Supplementary Figure 1). Descriptively, similar to Pool et al.[1], participants with low levels on the measure that corresponds with affective stress were more likely to demonstrate habit formation following extensive training whereas those with higher levels of this measure tended to respond habitually already after short training (Supplementary Figure 1), but notably, our results did not reach statistical significance.

We also ran similar tests in which we simply integrated either the STAI score or the TICS chronic worrying subscale score instead of the stress affect factor in the above-mentioned mixed-effects regression model. We did not find any three-way interaction with devaluation and group for either of these scores (ps>0.11).

### Supplementary figures

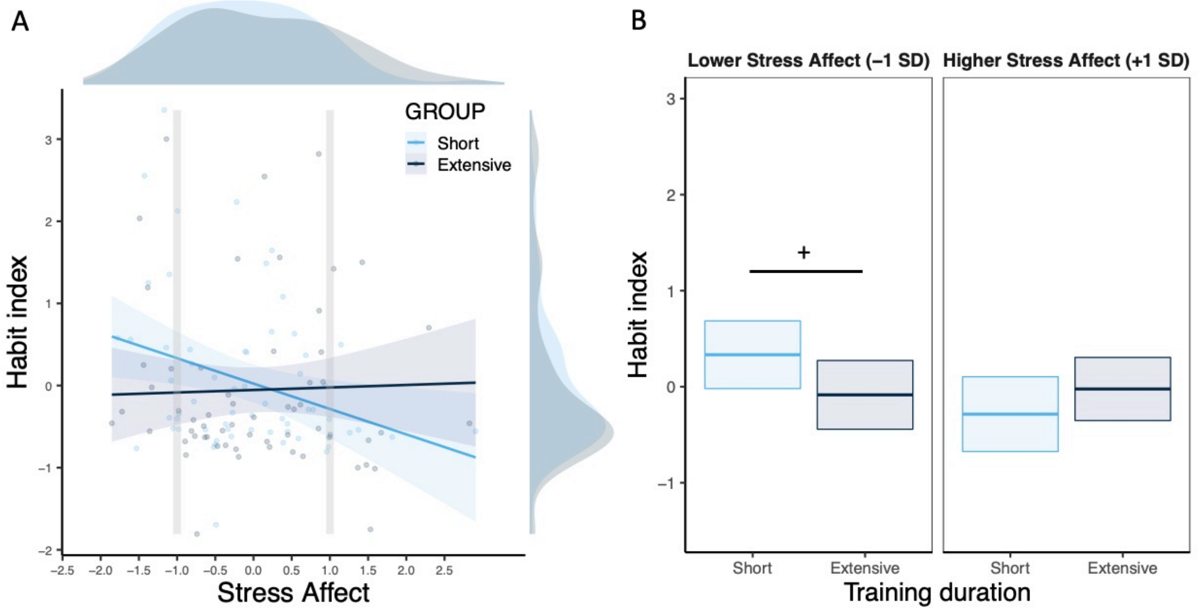


**Supplementary Figure 1.** (A) Habit index as a function of stress affect. The shaded areas represent 95% confidence intervals. (B) Estimated marginal habit index means for participants low (-1 SD) and high (+1 SD) on the stress affect measure in each experimental group. The boxes represent the area between 95% confidence intervals of the means.

* For the habit index higher indices represent more goal-directedness and values around zero represent habitual responders. The stress affect factor was inferred from exploratory factor analysis.

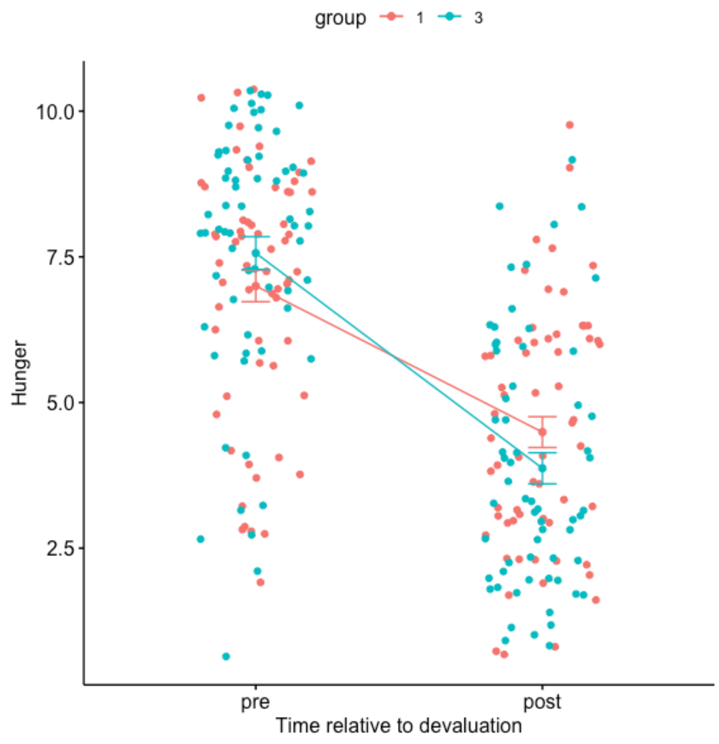


**Supplementary Figure 2.** Hunger ratings before and after outcome devaluation for the (remained) valued and devalued outcomes in each group.

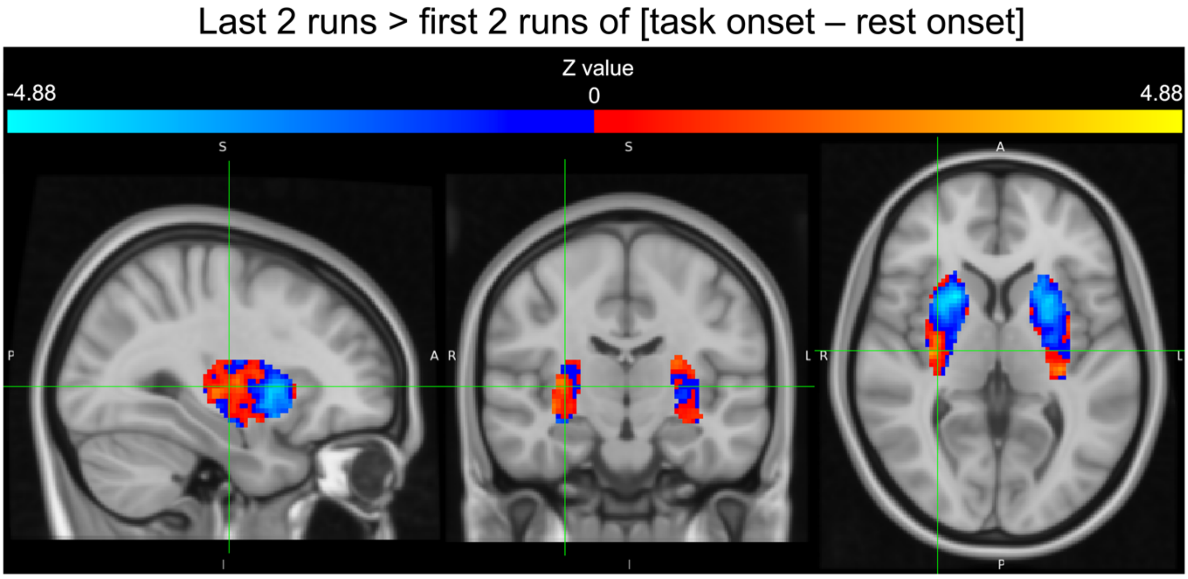


**Supplementary Figure 3. Differential training-related BOLD response patterns across the bilateral putamen.** The anterior and posterior parts of the bilateral putamen show clear distinct response profiles. The anterior parts reduced their activations to task cues following extensive training whereas the posterior parts appear to increase their activity (although not to the same extent). The statistical maps show **unthresholded Z-score statistics** of task cue onsets vs. rest cue onsets following extensive training (last 2 runs vs. first 2 runs) extracted from the mask used for the small volume correction analyses in the bilateral putamen. Cold and hot colors indicate decrease and increase in activations toward task cues following extensive training, respectively.

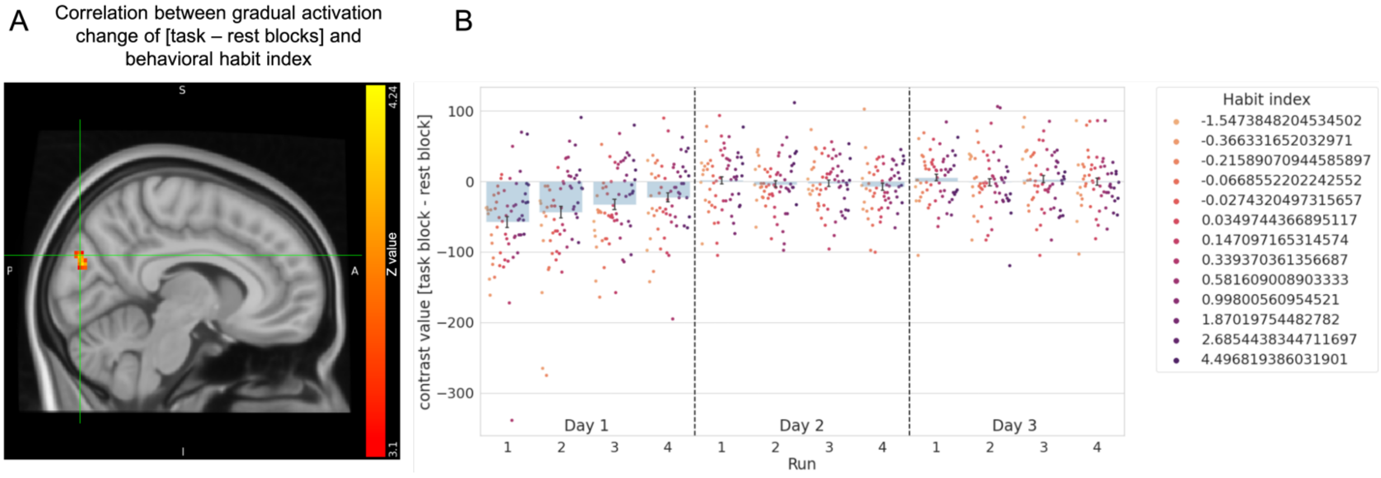


**Supplementary Figure 4**. **Correlation between the behavioral habit index and a linear trend in training duration-induced changes in neural activity (results from the secondary parallel analysis of the entire blocks). (A)** Whole-brain analysis correlating linear trends in activation across training of [task blocks vs. rest blocks] with the individual habit index revealed a negative correlation with goal-directedness in the right cuneal cortex. **(B)** Plots are presented for illustrative purposes of the effects in **panel A:** Averaged contrast values of task vs. rest blocks within the identified significant cluster in the right cuneal cortex shows that this effect is driven by more activation in this region during rest compared with task blocks in early stages of the task whereas it became indifferent in later stages in habitual participants but not in goal-directed participants where it was indifferent throughout the entire task.

Notes: (1) The fMRI results were cluster-corrected ($p<0.001$) as a strict threshold to account for multiple comparisons in this exploratory parallel analysis. (2) The habit index is structured such that higher values indicate goal-directed behavior and values around 0 indicate habitual responding. (3) Error bars in bar plots indicate 68% confidence interval (equivalent to ±1 std error).

### Supplementary tables

**Supplementary Table 1. Factor loadings obtained by exploratory factorial analysis of the subscales of the TICS, BIS-11 and STAI questionnaires.**

| | Factor A | Factor B - **labeled Stress Affect** | Factor C |
|---|---|---|---|
| TICS Social overload | **0.825** | -0.165 | 0.043 |
| TICS pressure to perform | **0.775** | 0.098 | -0.102 |
| TICS work discontent | **0.462** | 0.376 | 0.132 |
| TICS excessive demands at work | **0.512** | 0.364 | 0.108 |
| TICS lack of social recognition | **0.721** | 0.012 | -0.006 |
| TICS social tensions | **0.548** | 0.141 | 0.113 |
| TICS social isolation | 0.070 | **0.714** | -0.167 |
| TICS chronic worrying | 0.147 | **0.768** | 0.043 |
| TICS work overload | **0.650** | 0.072 | -0.020 |
| BIS motor | 0.165 | -0.134 | **0.719** |
| BIS attentional | 0.000 | 0.426 | **0.515** |
| BIS nonplanning | -0.092 | 0.017 | **0.831** |
| STAI state-trait (combined) | -0.043 | **0.883** | 0.038 |

The larger loading for each subscale is highlighted in bold.

**Supplementary Table 2. Cluster activation table of the (positive) correlation between valued vs. devalued (whole) blocks and the behavioral habit index across all participants.**

| Cluster Index | Voxels | P | -log10(P) | Z-MAX | Z-MAX X (mm) | Z-MAX Y (mm) | Z-MAX Z (mm) | Z-COG X (mm) | Z-COG Y (mm) | Z-COG Z (mm) | COPE-MAX | COPE-MAX X (mm) | COPE-MAX Y (mm) | COPE-MAX Z (mm) | COPE-MEAN |
|---|---|---|---|---|---|---|---|---|---|---|---|---|---|---|---|
| 11 | 2132 | 1.950000e-38 | 37.70 | 8.69 | -36 | -30 | 63.6 | -36.00 | -27.00 | 63.10 | 83.30 | -30 | -24 | 75.6 | 23.70 |
| 10 | 909 | 2.480000e-21 | 20.60 | 6.85 | 6 | -64 | -13.2 | 13.20 | -57.40 | -20.20 | 18.50 | 2 | -68 | -13.2 | 9.77 |
| 9 | 352 | 7.440000e-11 | 10.10 | 5.05 | -14 | -2 | 46.8 | -6.75 | -14.80 | 49.40 | 20.70 | 0 | -8 | 66.0 | 10.00 |
| 8 | 304 | 9.490000e-10 | 9.02 | 4.74 | -42 | -78 | 6.0 | -46.70 | -74.50 | 5.79 | 25.00 | -56 | -74 | 1.2 | 14.80 |
| 7 | 298 | 1.320000e-09 | 8.88 | 5.26 | 38 | -12 | 54.0 | 37.20 | -22.30 | 58.30 | 24.40 | 34 | -26 | 73.2 | 12.60 |
| 6 | 253 | 1.650000e-08 | 7.78 | 5.18 | -58 | -20 | 22.8 | -49.50 | -25.00 | 21.70 | 22.70 | -58 | -20 | 22.8 | 11.90 |
| 5 | 130 | 4.300000e-05 | 4.37 | 5.18 | 42 | -66 | 6.0 | 46.10 | -69.40 | 4.84 | 21.00 | 48 | -70 | 3.6 | 13.00 |
| 4 | 101 | 3.810000e-04 | 3.42 | 4.95 | -12 | -48 | -15.6 | -15.00 | -50.70 | -16.80 | 10.40 | -16 | -58 | -13.2 | 6.74 |
| 3 | 89 | 9.960000e-04 | 3.00 | 4.44 | 40 | -32 | 20.4 | 40.70 | -28.20 | 20.50 | 15.10 | 40 | -32 | 18.0 | 9.60 |
| 2 | 80 | 2.110000e-03 | 2.68 | 4.97 | -14 | -24 | 6.0 | -13.30 | -23.40 | 6.68 | 9.71 | -4 | -24 | 8.4 | 6.81 |
| 1 | 51 | 2.910000e-02 | 1.54 | 4.91 | -64 | 2 | 32.4 | -61.70 | 2.13 | 33.20 | 30.80 | -64 | 2 | 32.4 | 17.70 |
